# MONET: A model for neutral evolution of traits

**DOI:** 10.64898/2026.04.16.718874

**Authors:** Isabela do O, Oscar Gaggiotti, Marianne Bachmann Salvy, Jerome Goudet, Pierre de Villemereuil

**Author notes:** Corresponding author: Isabela do O, Department of Ecology and Evolution, University of Lausanne, Lausanne, Vaud 1004, Switzerland. Equally contributing authors (Jerome Goudet, Pierre de Villemereuil).

## Abstract

Demonstrating that local adaptation drives phenotypic divergence in quantitative traits requires distinguishing selection from neutral differentiation. Existing methods for detecting selection on quantitative traits, particularly *Q*_*ST*_ – *F*_*ST*_ comparisons, rely on simplified assumptions about population structure. These methods assume equal relatedness among all subpopulations, which is not necessarily true in a real metapopulation. When assumptions are violated, it has been shown that *Q*_*ST*_ – *F*_*ST*_ leads to elevated false positive rates. Here we present MONET, a model for the neutral evolution of traits that creates a robust baseline against which hypotheses on trait evolution can be tested. We show how the null model provided by MONET can be the basis of two tests: one based on the log-ratio of ancestral variances (log_*AV*_) statistics we previously introduced, and another, based on testing the influence of environmental variables from the populations of origin on the phenotypic traits measured in a common garden. MONET is implemented in an R-package as a Bayesian linear mixed-effect framework modeling population structure through relatedness matrices. We show through extensive simulations across different population structures and selective scenarios how our two hypothesis tests based on MONET maintain proper calibration while maintaining or exceeding the power of alternative methods.

## Introduction

Understanding how quantitative traits evolve in spatially structured populations requires distinguishing the contributions of several processes acting simultaneously, most notably genetic drift, gene flow, and natural selection. In any finite metapopulation, subpopulations will diverge through the stochastic accumulation of allelic differences, even in the complete absence of selection (Lynch 1994; Lynch *et al*. 1998). Any inference about the evolutionary forces shaping trait variation, whether testing for local adaptation, identifying environmental drivers of divergence, or evaluating the role of sex-specific selection, depends on a clear expectation of what trait variation should be under neutrality alone. A neutral model of trait evolution accounting for the history of genetic drift and migration between populations is thus an essential requirement for any evolutionary study aimed at elucidating the mechanisms that generate phenotypic diversity.

Wright (1949) emphasized the importance of genetic drift as a driver of evolutionary change and developed a theory for evolution in finite populations that partitions the molecular diversity into within-population and between-population components, allowing for a measure of divergence between subpopulations, *F*_*ST*_, which remains the standard measure of the proportion of genetic variation attributable to population structure. Based on this theory, Kimura (Kimura and others 1968) asserted that the vast majority of evolutionary changes observed at the molecular level are not the result of selection but rather the random fixation of selectively neutral or nearly neutral mutations through genetic drift, and was the first to propose that a model of neutral evolution should serve as the null hypothesis against which evidence for selection can be evaluated. From this premise, the theory derives quantitative predictions: the rate of molecular evolution is approximately equal to the neutral mutation rate (Kimura 1983), and the expected heterozygosity within a population scales with 4*N*_*e*_*µ* (where *N*_*e*_ is the effective population size, and *µ* the mutation rate) (Watterson 1975).

At the quantitative trait level, differentiation is described by additive genetic variance rather than allele frequencies. The general expectation under neutrality is that the additive genetic variance between subpopulations scales with *F*_*ST*_, specifically that *V*_*B*_ = 2*F*_*ST*_*V*_*A*_, where *V*_*B*_ is the additive genetic variance between subpopulationas, and *V*_*A*_ is the ancestral additive genetic variance (Cockerham and Tachida 1987; Lande 1992; Whitlock 1999). This relationship provides the general reference point connecting molecular estimates of population structure to expectations for quantitative trait divergence.

However, *V*_*B*_ = 2*F*_*ST*_*V*_*A*_ relies on all pairs of subpopulations to be equally related so that a single scalar *F*_*ST*_ averaged across all pairs fully describes the coancestry structure of the metapopulation. This equality comes from deriving the expected between-subpopulation additive variance directly from drift and mutation (Lande 1992): 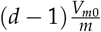, where *V*_*m*0_ is the mutational variance, *m* is the migration rate, and *d* is the number of subpopulations. At drift-migration equilibrium, this island model also yields 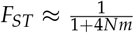, and substituting this expression recovers 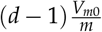 as an instance of *V*_*B*_ = 2*F*_*ST*_*V*_*A*_.

In topologies where relatedness is not isotropic, a scalar *F*_*ST*_ does not provide a link between the molecular and the additive genetic variance between subpopulations. For example, under a stepping-stone model, pairwise relatedness decays with geographic distance, so neighboring subpopulations are more similar than distant ones even when their average *F*_*ST*_ is the same (Kimura and Weiss 1964). Under a hierarchical population structure, relatedness will differ among within-cluster and between-cluster pairs of subpopulations (Slatkin and Voelm 1991). In both cases, a single averaged value of *F*_*ST*_ is compatible with many different underlying patterns of pairwise relatedness, and so no longer determines a unique expectation for *V*_*B*_.

To address these challenges in the context of detecting signs of local adaptation, do O *et al*. (2025) developed an approach based on a new test statistics named *log* _*V*_ . The approach expands the equation *V*_*B*_ = 2*F*_*ST*_*V*_*A*_ and *V*_*W*_ = (1 − *F*_*ST*_)*V*_*A*_ (Whit- lock 1999) to account for the variance-covariance pattern of trait distribution by substituting *F*_*ST*_ with a coancestry matrix that describes the full pattern of pairwise relatedness. The *log*_*AV*_ test statistics is then computed as a log-ratio of estimators of the same ancestral additive genetic variance, one based on within-population variation, the other based on between-population variation. Here we expand on this approach to formalize a general model of neutral evolution of traits (MONET), which allows for a series of hypothesis tests, including *log* _*AV*_. We show the usage of the neutral model for detecting the impact of environmental variables on trait distributions and how it can be used for understanding ecological processes such as invasion patterns by reanalyzing published datasets.

The aims of the present study are fourfold: (i) to present the MONET framework: a Model for Neutral Evolution of Traits; to evaluate the calibration and statistical power through extensive simulations, benchmarking MONET and two widely used methods for identifying local adaptation, *Q*_*ST*_ –*F*_*ST*_ (Whitlock and Guillaume 2009) and Driftsel (Ovaskainen *et al*. 2011; Karhunen *et al*. 2013); (iii) to illustrate, using a case study in which the population structure deviates from the island model, how different neutral models account for such structure (Ramirez-Valiente *et al*. 2022); and (iv) to demonstrate how MONET can be used to formally address complex biological questions, using a case study in invasion ecology (Shirk and Hamrick 2014).

## Method description

MONET implements a Bayesian linear mixed-effects model of neutral trait evolution in spatially structured populations. The model describes the expected distribution of trait values under the distribution of relatedness generated by forces such as drift, gene flow, and the demographic history of the metapopulation, providing an explicit null model against which deviations can be tested. From this neutral baseline, several hypothesis tests follow naturally: the log_*AV*_ test (do O *et al*. 2025) for local or global adaptation (Yeaman 2022), tests for environmental associations through fixed-effect covariates, and tests for other biological effects (effects of sex, age, plasticity, etc.), including their interaction with adaptation. The framework was designed for quantitative traits measured in common gardens, where environmental variance is standardized, and the observed variation can be interpreted as reflecting heritable differences shaped by the metapopulation’s demographic history.

The model estimates ancestral additive genetic variance by decomposing the demographic history into two aspects, one that describes the coancestries between subpopulations (*V*_*A,B*_), and another that describes the coancestries between individuals while correcting of the shared history coming from the subpopulation level (*V*_A,*W*_). Under neutral evolution, both estimates correspond to the same ancestral variance partitioned through different aspects of the demographic history captured by population structure (Lynch *et al*. 1998; Whitlock 1999), and are therefore expected to be equal on average.

Keeping the two variances separate, rather than collapsing them into a single combined relatedness matrix (as in Eq. 12 in (Ovaskainen *et al*. 2011)), is a deliberate design choice. A unified formulation would absorb any unexplained signal, such as local adaptation on a variable not included in the model, into the unique ancestral additive genetic variance estimate, or worse, into the fixed effects estimates, resulting in a biased estimation and an inflated false positive rate. The separation allows the model to reveal mismatches between the two sources of information (within and between subpopulations), without forcing an improper fit. From this design, two complementary tests arise: a test for the equality between the estimated ancestral variances *V* _*A,B*_ and *V* _A,W_ through the log-ratio, and tests on fixed-effect coefficients describing environmental or other covariates.

The neutral model is formalized by the following Bayesian linear mixed-effects model, implemented using the R package `brms` (Bürkner 2018):

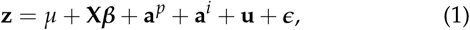

where **z** is the trait value, *µ* is the overall phenotypic mean, **a**^*p*^ is the population-level random effect, **a**^*i*^ is the individual-level random effect, and **ffl** is the residual error. The notation **u** represents additional random effects (e.g., common garden blocks, years), which may be added to the model. **X** is the design matrix relating individual measures to the vector of fixed effects ***β***. The matrix **X** can include individual-specific characteristics (e.g., sex, age, body size), or experimental design factors (e.g., common garden environmental variables, time of measurement). **X** can also be used to describe environmental covariates; for each individual, the covariate value corresponds to the environmental condition of their population of origin. We show below how to use this to construct a statistical test for local adaptation. Continuous variables are standardized (mean-centered and scaled by standard deviation) in MONET prior to model fitting to improve MCMC (Markov chain Monte Carlo) sampling efficiency. Note that this formulation generalizes the original log_*AV*_ formulation (see Eq.8 in do O *et al*. (2025)) because it includes additional random and fixed effects.

The between- and within-population additive genetic effects, **a**^*p*^ and **a**^*i*^, are randomly distributed and follow multivariate normal distributions:

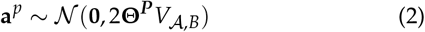

and

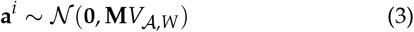

where *V*_*A,B*_ is the ancestral additive genetic variance defined through between-population effects and *V*_A_ _,*W*_ is the ancestral variance defined through within-population effects. Under neutrality, we expect *V*_*A,B*_ = *V*_*A,W*_ because both correspond to the same ancestral variance, but use different aspects of the demographic history, which are modeled through relatedness.

Briefly, **Θ**^*p*^ is the population-level relatedness matrix, an *r × r* symmetric matrix, where *r* is the number of subpopulations. Elements 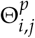 represent the average probability of identity by descent (IBD) between a random pair of alleles, one sampled from subpopulation *i* and one from subpopulation *j*, relative to the ancestral population. Diagonal elements 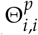 represent the average relatedness for pairs of alleles sampled from different individuals within subpopulation *i* (do O *et al*. 2025).

The individual-level relatedness matrix **M** is an *n × n* matrix describing genetic relationships among F1 individuals within subpopulations, after accounting for the shared ancestry already captured in **Θ**^*p*^. Usually, in common garden studies, breeding occurs only within populations (no cross-population matings), making **M** a block-diagonal matrix, with blocks corresponding to each subpopulation; see do O *et al*. (2025). **M** can be calculated using genetic or pedigree data. Note that we use F1 here for the sake of simplicity, but MONET can accommodate complex and deeper crossing designs, even if they include multiple generation steps (e.g., F2) from the parental generation.

We use default priors for fixed and random effects, which can be adapted to the researcher’s study system. The Bayesian framework yields full posterior distributions for all parameters, including 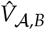 and 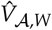, naturally incorporating parameter uncertainty. (Detailed model specification, parameters, and convergence diagnostics are provided in the MONET package documentation and vignette (do O *et al*. 2026).)

### Hypothesis tests from the neutral model

#### Patterns of divergence that deviate from neutrality

Several hypothesis tests follow directly from the neutral model. The most obvious, and most analogous to existing methods for detecting selection, compares the two estimates of ancestral genetic variance via their log-ratio, the log_*AV*_ test statistic:

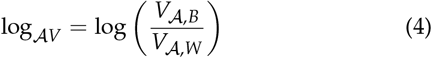

Under neutrality, E [log_*AV*_] = 0. A log-ratio significantly greater than zero indicates local adaptation (*V* _*A,B*_ > *V*_A_ _,*W*_), suggesting that phenotypic divergence among populations exceeds neutral expectations. Conversely, a log-ratio significantly less than zero indicates spatially-uniform selection (*V*_*A,B*_ < *V*_*A,W*_), suggesting that selection has reduced phenotypic divergence among populations relative to what would be expected under drift alone. We interpret the former scenario as local adaptation and the latter as global adaptation. Here we refer to global adaptation according to Yeaman (2022), meaning that all analyzed subpopulations are exposed to an identical fitness landscape with the same optimum phenotype, which would yield lower phenotypic divergence than expected by drift, i.e., more similarity between populations than expected.

From the posterior distributions of *V*_*A,B*_ and *V*_*A,W*_, we calculate the log-ratio for each posterior sample and compute a Bayesian *p*-value as:

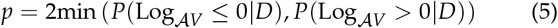

This *p*-value quantifies the proportion of the posterior where the log-ratio has the opposite sign from the posterior median (used here as the point estimate), doubling this proportion for a two-tailed test. This Bayesian *p*-value has been shown to have asymptotic properties similar to frequentist *p*-values (Shi and Yin 2021), which we validate here by checking the calibration of MONET.

#### Correlation between divergence and covariates

A second class of tests evaluates fixed-effects coefficients in the model. When environmental data, for example, are included as covariates in **X**, the corresponding regression coefficients *β* quantify the association between trait values and environmental differences across populations, while controlling for neutral structure through the coancestry matrices in the random effects defined in Eq. 2 and 3. The same idea applies to any other covariate relevant to the problem (age, sex, or experimental factors), each resulting in a test of whether that covariate explains part of the variation beyond what the neutral model predicts. The test uses the same Bayesian *p*-value calculation as the log-ratio test in (Eq. 5), evaluating whether each *β*_*j*_ differs significantly from zero:

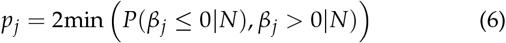

A significant environmental coefficient indicates that trait values (measured in a common garden environment) are systematically associated with that environmental variable across populations, even while accounting for neutral population structure.

### Using MONET

A step-by-step description of how to apply the model in the case of testing for local adaptation is present in Figure 3 of do O *et al*. (2025).

MONET requires three primary types of data:

1. **Genetic data from the parental generation:** SNPs, microsatellites, or other neutral markers from individuals sampled from each subpopulation in the wild. This data is used to estimate the population-level relatedness matrix **Θ**^*p*^, which captures the demographic history and relatedness structure among subpopulations. The MONET R package expects genotype data in dosage format, where genotypes are encoded as 0, 1, or 2, representing the number of copies of the alternative allele. If this data is unavailable, parental genotype may be reconstructed from F1 genetic data using software such as Sequoia (Huisman 2017) or COLONY (Jones and Wang 2010).
2. **Phenotypic measurements from the offspring generation (F1):** Quantitative trait values from individuals reared in a common garden environment to control for environmental effects.
3. **Relatedness information for F1 individuals:** Either (a) genetic data from the F1 generation to estimate relatedness between individuals, or (b) pedigree information.

Again, we use F1 for the sake of simplicity, but MONET can accommodate more complex and deeper crossing designs. A complete user manual and tutorial are available as vignettes of the R package.

## Method testing

Although MONET’s broader value lies in providing a flexible neutral model for trait evolution, evaluating its statistical properties is most naturally done through the hypothesis tests it allows for. Here we focus on two tests, the log_*AV*_ test and the environmental covariate test, and benchmark them against two widely-used methods for detecting selection: Whitlock and Guillaume (2009) implementation of *Q*_*ST*_ – *F*_*ST*_ and Driftsel (Ovaskainen *et al*. 2011; Karhunen *et al*. 2013). These two methods were chosen given their vast usage and relevance in the literature, as they have been cited over 600 and 200 times, respectively. We provide a succinct description of both in the supplementary information section.

### Simulations

To test the calibration and power of MONET (and compare it to earlier methods), we simulated metapopulations using QuantiNemo2 (Neuenschwander *et al*. 2019) under different population structures (island model or IM, stepping stones or SS, and hierarchical; see Figure 1) with phenotypic traits evolving either neutrally or under various selective pressures. Specifically, the hierarchical structure refers to a situation in which an ancestral population splits into separate lineages without gene flow between lineages, with each lineage subsequently splitting into another set of lineages. This population structure, illustrated in the lower left panel of Figure 1, is based on previous works (Karhunen *et al*. 2013; do O *et al*. 2025). All simulated metapopulations described in this manuscript have a common ancestral population; the island and stepping stones models have ongoing gene flow between present subpopulations, while the hierarchical model does not have ongoing gene flow.

**Figure 1.**
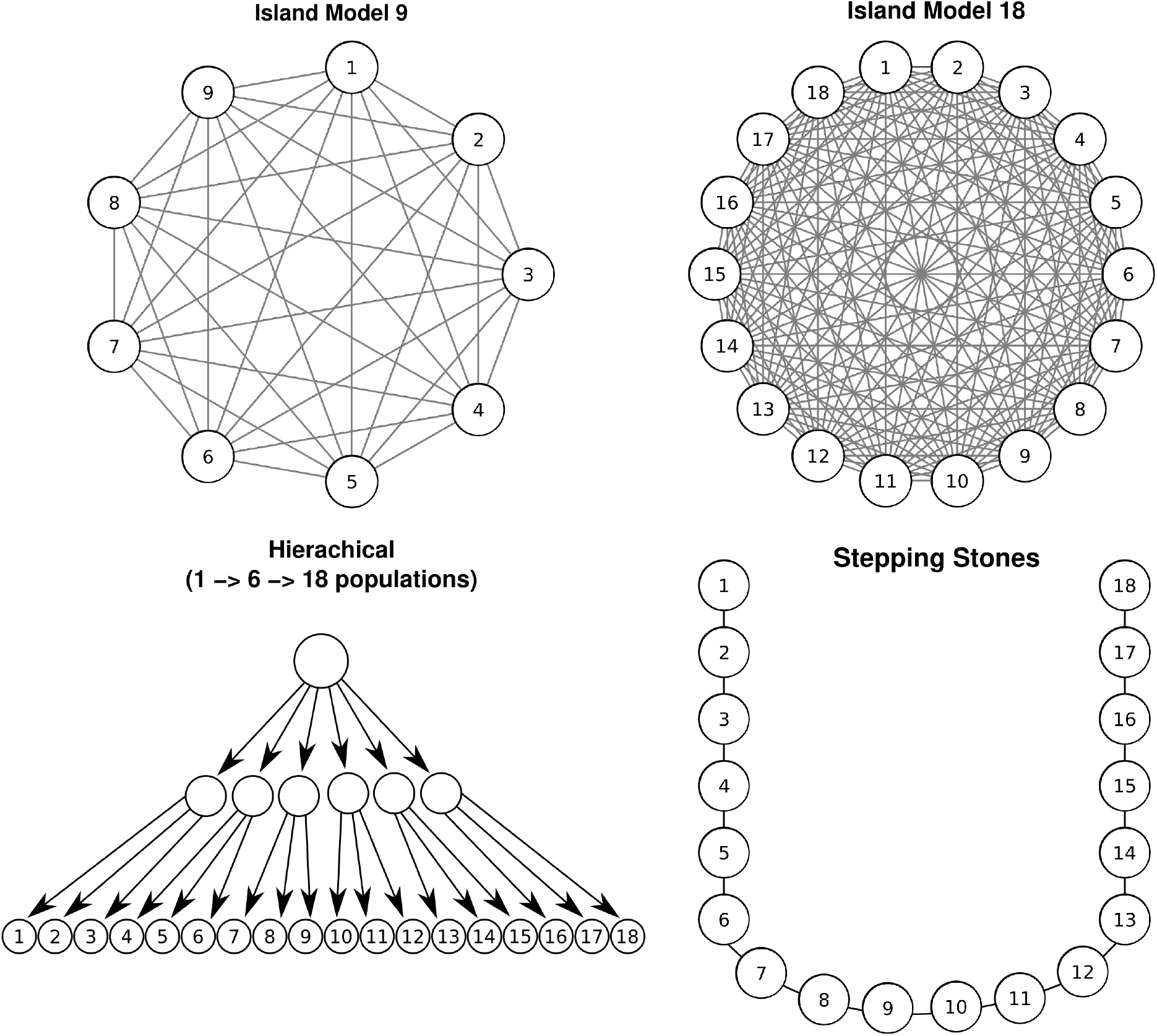
Simulated population structures with lines describing gene flow between populations and arrows showing ancestry relation.

We will refer to replicates with the same set of parameters as a “scenario”. To modulate the strength of selection, we varied two key parameters controlling selection strength (Figure 2). First, we varied the width of the peak in fitness 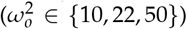, which is inversely related to the strength of within-population stabilizing selection. Second, we varied the distance between environmental optima, denoted Δ*θ*, which represents the difference between the most divergent environments (Δ*θ* ∈ {0, 4.6, 10}) and is related to the strength of between-population local adaptation. Note that for Δ*θ* = 0, the selection optimum is the same across all populations, leading to phenotypic and genetic differentiation below those expected under neutrality, i.e. the opposite of local adaptation. As mentioned above, and following Yeaman (2022), we refer to such a case as ‘global adaptation’.

**Figure 2.**
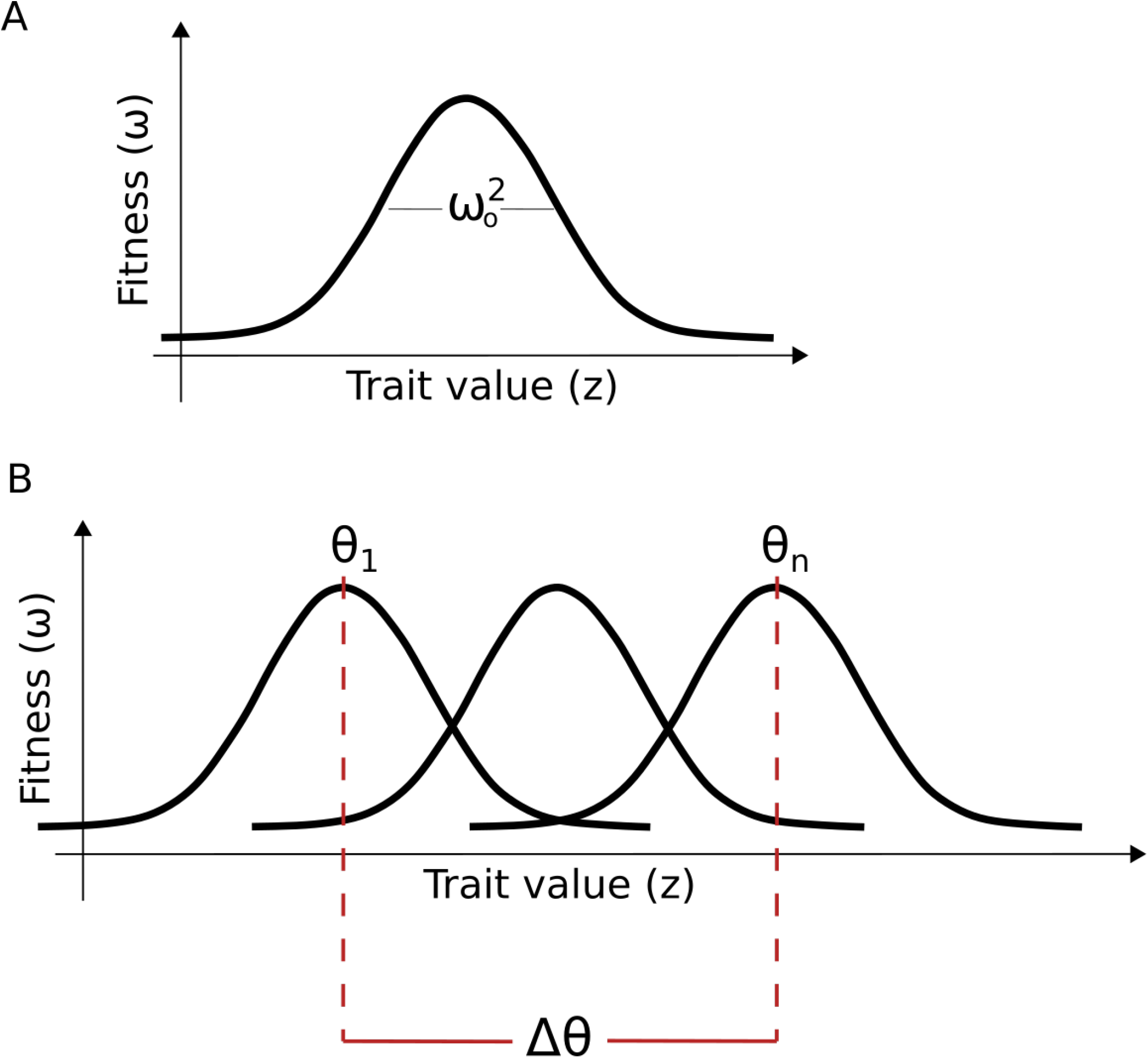
Visual description of the two parameters used to describe the selection that the metapopulations experienced. (A) shows the width of the fitness function, 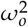. (B) shows the distance between the optima of the most distant environments, Δ*θ*.

For stepping stones and hierarchical models, we additionally considered the correlation patterns between environmental similarity and demographic connectivity or history (Figure 3). The parameter that we will hereinafter refer to as “correlation pattern” describes whether the neutral and the selective divergence follow the same pattern, i.e if environmental differences increase with genetic distances. For stepping stones, we simulated two types of correlations: parabola and cline. In the cline, the further apart two populations are, the more their optima differ, while in a parabola, the monotonic decrease in environmental correlation with distance is broken, such that populations far apart can experience similar environmental conditions. In the hierarchical model, we exposed metapopulations to two types of correlations that we named “grouped” and “swapped”. Under the grouped case, subpopulations within the same clade (i.e., more closely related) are exposed to the same selective pressures. Under the swapped case, subpopulations within a clade are exposed to different pressures.

**Figure 3.**
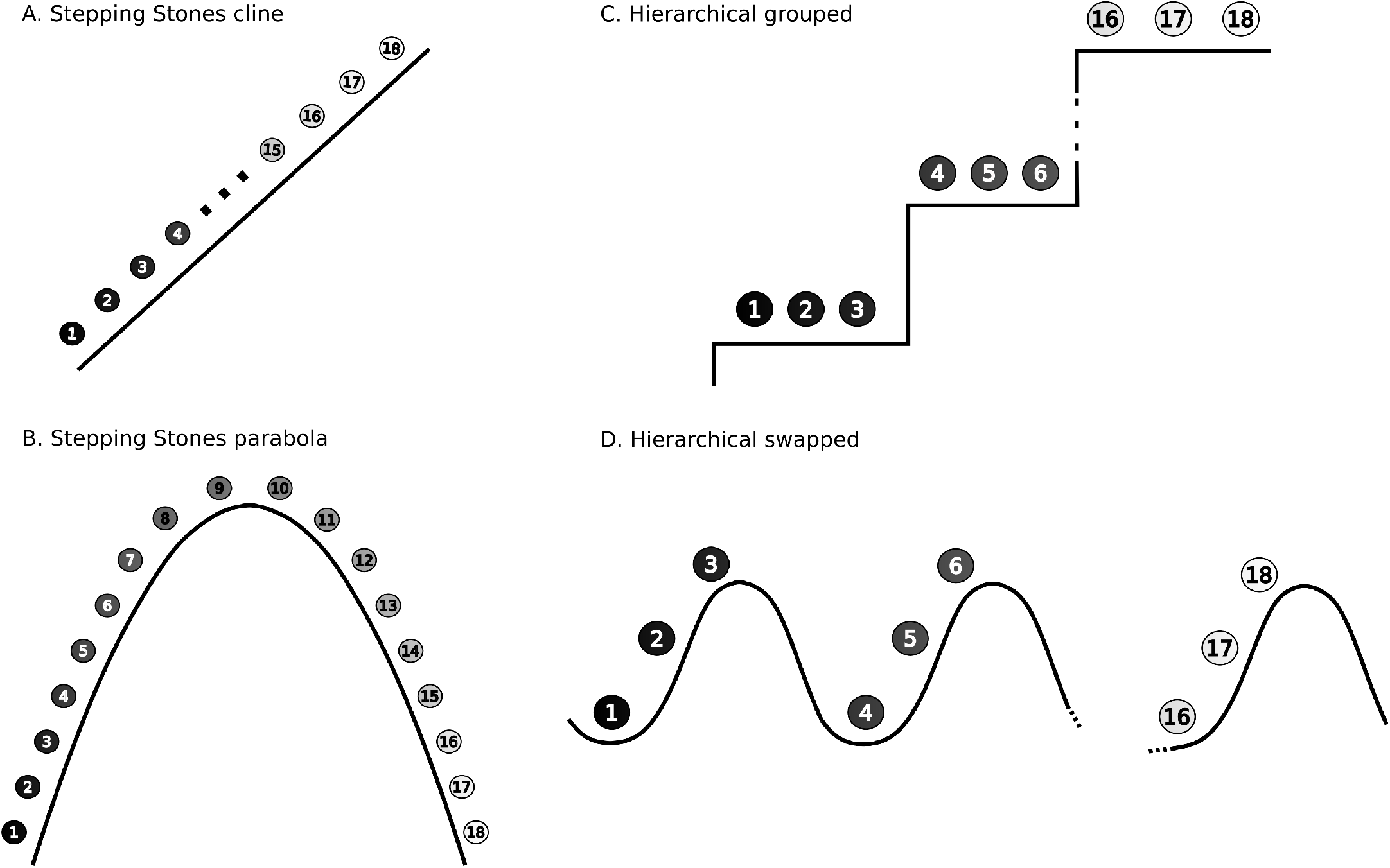
Graphical description of the types of correlation patterns between the neutral and the adaptive divergence. Shading describes the similarity in relatedness between pairs of subpopulations, i.e., two circles of similar shade are more related. The x-axis indicates the different population IDs, while the y-axis indicates the optimum trait value for each subpopulation. A. The “cline” correlation type, applied to stepping stones population structure, subpopulations more closely related are subject to more similar environments. B. The “parabola” correlation type, applied to stepping stones population structure, subpopulations at the edge of the stepping stones are exposed to more similar environments, and subpopulations in the center are exposed to more different environments. C. The “grouped” correlation type, applied to the hierarchical structure, populations from the same clade experience the same environment. D. The “swapped” correlation type, applied to hierarchical structures, populations from the same clade are exposed to different environments.

We generated 500 replicates per scenario. For each of them, we sampled 10 individuals per subpopulation and characterized their neutral genome (2000 bi-allelic loci) and a quantitative trait. Specifically, we considered two quantitative traits: (1) a trait under selection, and (2) a neutral trait. Both traits were determined by 100 additive bi-allelic loci, with uniformly distributed allelic effect, and a mutation rate of 10^−7^ and free recombination between loci. To verify that selection did not affect neutral population structure, we confirmed that neutral traits from scenarios with selection were statistically indistinguishable from those in scenarios without selection.

For all three structures (island model, stepping stones, and hierarchical), there were 18 subpopulations, and we also simulated the island model with only 9 subpopulations as shown in Figure 1.

For testing the implementation of environmental covariates in MONET, we refer throughout the current manuscript as “MONET w/environment”. The “environment” information we used in our tests was the optimum value assigned to each subpopulation in our simulations.

#### Demographic and selection parameters

We first evaluated how population-level parameters affect the performance of each method. These parameters included selection strength, environment differentiation, and demographic structure as previously described. For these, we kept the same “common garden” design where we sampled 10 individuals per population per replicate and established a North Carolina Design II breeding scheme (as described by Beavis *et al*. (2023)), in which every individual of one sex (5 in this case) mates with every individual of the other sex (5). Each cross produced two offspring characterized by their quantitative trait values and neutral genomes. We applied all three methods (Driftsel, MONET, and *Q*_*ST*_ –*F*_*ST*_) to identical sets of individuals and trait values within each replicate.

#### Sampling design and common garden parameters

Besides population-level parameters, we compared how sampling strategies and common garden designs influence method performance. Specifically, we tested the impact of: 1. the number of sampled subpopulations (9 vs. 18), 2. the number of F1 individuals analyzed per subpopulation (25 vs. 50), and 3. the crossing design (1 *×* 9, 3 *×* 3, or 5 *×* 5 matings per subpopulation, where the first value can be interpreted as the number of dams sampled in the population and the second as the number of sires). With these comparisons, we assessed how experimental design choices affect the statistical power of the three methods. For the first parameter, the number of sampled subpopulations, we sampled 9 populations from the middle of the stepping stones structure (subpopulations 6 to 14 on Figure 1). For hierarchical, we picked two monophyletic clades (subpopulations 4 to 12). As for the island model, we did not subsample; instead, we simulated metapopulations with 18 and 9 subpopulations. The down-sampling of the number of individuals in the F1 generation was done by randomly sampling 25 individuals per subpopulation from the 25 families, instead of analyzing all 50 individuals. For the mating design, in the 3 *×* 3 case, we sampled 6 individuals instead of 10 and crossed them following the North Carolina II breeding scheme (Beavis *et al*. 2023), again with 2 offspring per mating. As for the 1 *×* 9, we chose one individual at random to be one sex and all others to be the other, which would all have 2 offspring per mating. Therefore, 3 *×* 3 and 1 *×* 9 schemes consider the same number of individuals, but differ in their relatedness structure.

### Analyses

We compared the methods in two ways: comparing the false positive rates (FPR) at different thresholds to assess the calibration, and using a receiver operating characteristic (ROC) curve for analyzing power.

#### Calibration

We evaluated method calibration by examining the relationship between nominal significance threshold values and observed FPR under neutral evolution across all population structures. We calculated these two measures for neutrally evolving traits in scenarios where there was local or global adaptation for another trait. Selection was not directly acting on any of the traits considered in this analysis, but the presence of selection in another trait could potentially impact the behaviour of the methods, which is why we performed this analysis in all selection scenarios.

Importantly, the “threshold” plotted on the x-axis of these calibration curves is not the same quantity for all three methods, and this changes the reference line that corresponds to good calibration. For *Q*_*ST*_ –*F*_*ST*_ and MONET, the x-axis is a nominal *p*-value threshold. Under the null, *p*-values are expected to be uniformly distributed. Thus, a well-calibrated method produces an FPR equal to the nominal threshold, i.e its threshold to FPR curve should fall on the one-to-one identity line. Curves lying above the diagonal indicate inflated FPR, and curves lying below indicate conservative behaviour.

For Driftsel, the x-axis is not a *p*-value but a sign of selection cutoff on the folded *S* scale. The *S*-statistic is bounded between 0 and 1, with values close to 0 or 1 indicating selection (Ovaskainen *et al*. 2011; Karhunen *et al*. 2013). Because both tails of *S* are interpreted as evidence of selection, we use the folded statistic *S*_folded_ = min(*S*, 1 − *S*) = 0.5− |*S* − 0.5| and plot 2(folded *S*) on the x-axis, which goes from 0 to 1. A replicate is normally considered significant when *S* < 0.2 or *S* > 0.8, adopting the criteria proposed by Ovaskainen *et al*. (2011). Thus, on the folded-*S*, it rejects neutrality when 2*×* folded_*S*_ falls below the cutoff, meaning when *S* lies within a distance of the cutoff divided by two of either 0 or 1, so given the adopted values of 0.2 and 0.8, it corresponds to 0.4. Since this cutoff does not follow the same distribution as a p-value, the threshold to FPR relationship is not expected to follow a one-to-one relationship. Instead, we assess Driftsel’s behaviour by asking whether its threshold-FPR relationship is stable across the different population structures.

#### Power

Switching to the analysis of the selected traits, to compare each method’s power to detect selection, we analyzed receiver operating characteristic (ROC) curves for each scenario. For MONET, this means evaluating the *log*_*AV*_ test and the environmental covariate test, both of which are applications of the underlying neutral model. ROC curves are a standard technique for visualizing diagnostic tests’ performance by plotting the true positive rate (sensitivity) against the false positive rate across decision thresholds (Fawcett 2004). This approach allows us to evaluate the discriminative ability of each method at a given FPR, independently of a particular significance threshold, making it particularly useful for comparing methods with different calibration properties. Specifically, comparing ROC curves allows for a fairer empirical comparison of power between methods that are not necessarily expected to be calibrated, such as *Q*_*ST*_ –*F*_*ST*_ in non-isotropic population structures, since the analysis is threshold-independent and focuses on the ranking of test statistics rather than their absolute values.

## Case studies

### A case study on Mediterranean oaks

To illustrate the use of MONET on empirical data, we reanalyzed the dataset of Ramirez-Valiente *et al*. (2022): this study aimed at understanding how climate, especially drought, has shaped differences in functional traits in metapopulations of two species of oaks, *Quercus faginea* and *Quercus lusitanica*. The authors measured a series of traits related to drought tolerance and growth in seedlings raised in common gardens, and asked whether population-level differences in these traits could be attributed to climate-driven selection. To test for selection, Ramirez-Valiente *et al*. (2022) applied both *Q*_*ST*_ –*F*_*ST*_ and Driftsel. To identify potential climatic drivers, they used H-tests (Karhunen *et al*. 2013) and Pearson correlations between trait means and climatic principal components.

This dataset was an attractive case study for two reasons. First, because the authors ran both *Q*_*ST*_ –*F*_*ST*_ and Driftsel on the same data, and we compared the usage of MONET for local adaptation analysis against the same two methods using simulated data. Second, the authors described patterns of population structure and indicated low neutral divergence and a history of postglacial expansion in the two species, suggesting a metapopulation structure that likely departs from the island model, an example of a situation in which MONET would be necessary.

Ramirez-Valiente *et al*. (2022) ran two common garden experiments, a greenhouse with two watering treatments (well-watered and dry) in both species, and an outdoor common garden for *Q. faginea* only. Here we are only reanalyzing the greenhouse experiment. They measured 11 quantitative traits in the greenhouse common gardens, these traits were: Specific leaf area (“SLA”), leaf lamina area (“area), leaf thickness (“thickness”), leaf perimeter-to-area ratio (“PA”), mass-based stomatal conductance (“gs.mass”), area-based stomatal conductance (“gs.area”), maximum quantum yield of photsystem II in light (“Fv’/Fm’”), Effective quantum yield of photosysmtem II (“ΦPSII”), non-photochemical quenching (“NPQ”), absolute height growth rate (“AGR”), relative height growth rate (“RGR”).

Phenotypic and genetic data, along with elevation and geographic coordinates for each population, were obtained from Ramirez-Valiente *et al*. (2022). Climatic principal components were recomputed from the same variables they used: the 19 WorldClim bioclimatic variables (BIO1 to BIO19) with the same resolution from 1970 to 2000 (Fick and Hijmans 2017). Soil pH (0–30 cm), and annual and summer indices of moisture were computed from monthly precipitation and potential evapotranspiration (Hargreaves and Samani 1985). For soil pH, we replaced the data from Trabucco and Zomer (2010) employed by Ramirez-Valiente *et al*. (2022) with SoilGrids 2.0 (Poggio *et al*. 2021), which is accessible from R and supports an easily reproducible pipeline. We followed Ramirez-Valiente *et al*. (2022) and combined the climate variables into principal components. Our first two PCs, similar to the original work, account for more than 70% of the variation (76.9% for *Q. faginea*, and 79.8% for *Q. lusitanica*.

To verify that our recomputed climate PCs reproduce the climatic ordering used by Ramirez-Valiente *et al*. (2022), given our substitution of the SoilGrids 2.0 soil pH layer, we compared per-subpopulation PC scores against those read off their Figure 3 for the analyzed subpopulations. Procrustes correlation on the PC_climate1_ and PC_climate2_ were high between ours and their PCs (*r* = 0.88, *p*-value = 0.001 for *Q. faginea*, and *r* = 0.90, *p*-value = 0.002 for *Q. lusitanica*). Per-axis Spearman correlations were strong for all axes, and cross-axis correlations were all near zero, confirming that our axes are not rotated into theirs. This step demonstrated that our substitution does not materially change the climate space analyzed.

First, we provide a visual comparison of the **Θ**^***P***^ matrix describing the metapopulation structure of *Quercus faginea* and *Quercus lusitanica* with the **Θ**^***P***^ of replicates of our simulated metapopulations. We then fit two MONET models to each of the two greenhouse datasets, each illustrating a different application of the model. The first model we will refer to as “base model” and only contains population and individual level coancestries as random effects, and it tests for signs of local or global adaptation only. The second one, which includes PC_climate1_ and PC_climate2_ as fixed effects, evaluates whether trait divergence tracks climate after accounting for neutral structure.

### A case study on the invasive species *Geranium carolinianum*

Beyond testing for associations with environmental covariates, MONET can be used to address a broader range of ecological questions. In this second case study, we illustrate its application to an invasion ecology question. Shirk and Hamrick (2014) compared phenotypic differentiation between native populations of *Geranium carolinianum* in the United States and invasive populations in China, asking whether trait divergence between the two ranges is consistent with adaptive evolution rather than genetic drift alone. Because a single *Q*_*ST*_ -*F*_*ST*_ comparison pooling all 20 subpopulations across the two countries would implicitly and wrongly assume the island model, the original analysis instead ran three separate *Q*_*ST*_ -*F*_*ST*_ tests per trait: one among the Chinese subpopulations, another for the U.S. subpopulations, and one treating each country as a single subpopulation, respectively testing for local adaptation within each range and for adaptive divergence between ranges. Here we show how a single MONET model can jointly address the same three questions. By including the coancestry matrices to capture the full nested population structure across both countries, together with country as a fixed effect, MONET simultaneously tests for local adaptation within each range through the log_*AV*_ statistic and for trait divergence between ranges through the country coefficient.

We ran MONET on the traits reported by (Shirk and Hamrick 2014), and included country as a covariate in our model, attempting to find whether divergence between the invasive and the native subpopulations was greater than that expected under neutrality. We obtained for each trait the log_*AV*_ value and its associated credible interval, along with the slope and the associated *p*–value for country.

For both case studies, we check for convergence and only report results for traits whose MONET model satisfied our convergence criteria (maximum 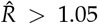 across all parameters, fewer than 1% divergent transitions, min ESS = 400 (Bürkner 2018)).

## Results

### Method calibration

Figure 4 shows the curves describing the false positive rates at different significance thresholds. Under the island model, *Q*_*ST*_ – *F*_*ST*_ and MONET were well calibrated (Figure 4, top three panels), consistent with the fact that this population structure satisfies the assumption of equal relatedness among subpopulations. However, once this assumption was broken (under the stepping stones and hierarchical scenarios), only MONET maintained proper calibration. This holds whether MONET’s test is based on log_*AV*_ or on the environmental covariate.

**Figure 4.**
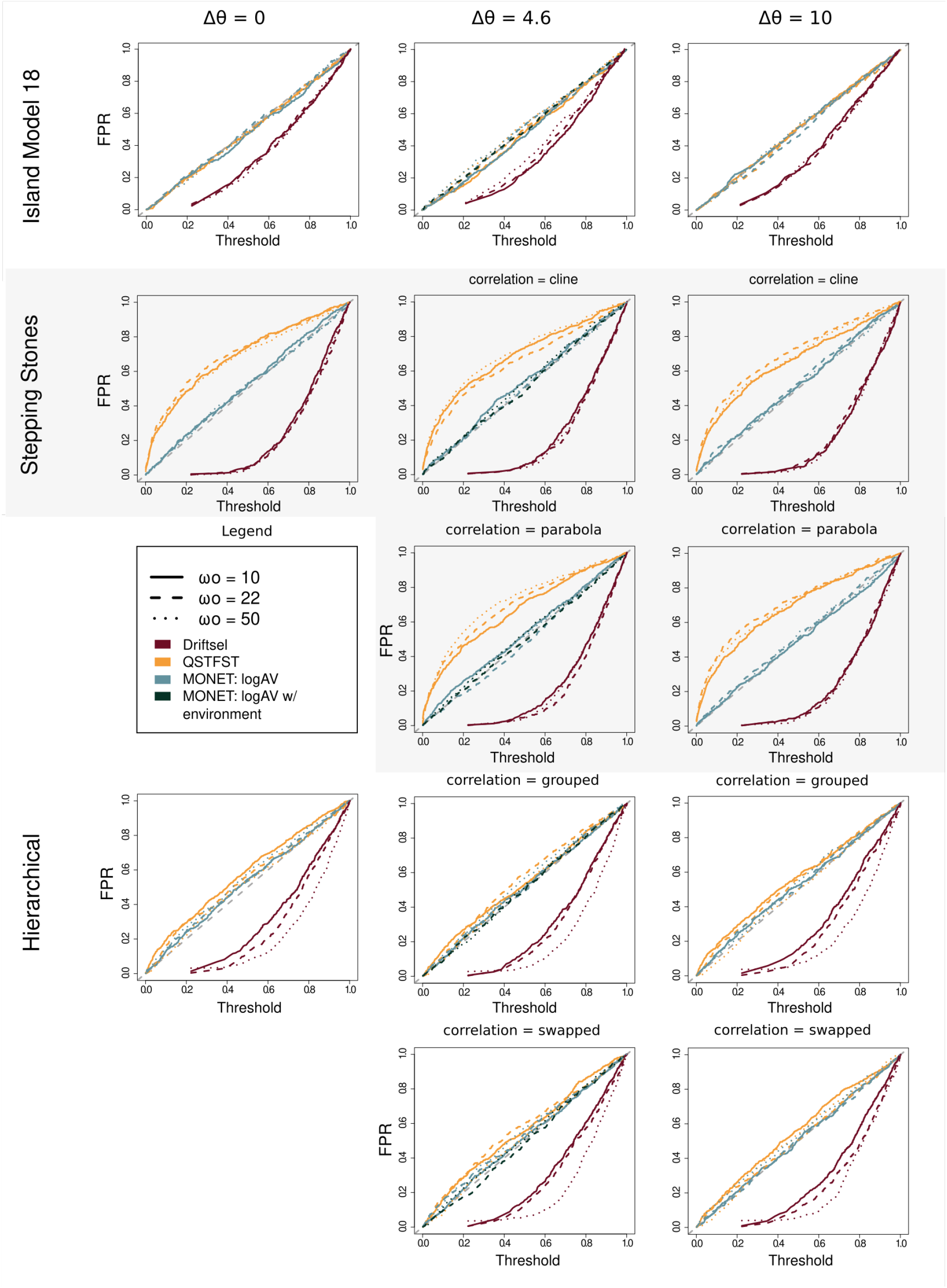
Tests calibration: Threshold by false positive rates (FPR) comparison for all three methods and all three population structures. Different strengths of selection are depicted by different line types, and colours show the different methods. The separate panels show the different scenarios related to the difference in the environment.

Under the stepping stones structure, *Q*_*ST*_ – *F*_*ST*_ showed systematic miscalibration (Figure 4, middle panels) across all analyzed scenarios within this structure. The threshold-FPR curve consistently fell below the diagonal, indicating an inflated FPR compared to the expected value from the nominal threshold. In contrast, MONET exhibited consistently calibrated threshold-FPR relationships across all stepping-stone scenarios. The contrast between *Q*_*ST*_ – *F*_*ST*_ and MONET was less pronounced in the hierarchical model, though *Q*_*ST*_ – *F*_*ST*_ still has a tendency to be miscalibrated (under the identity line, thus slightly inflated FPR).

As expected and described in the methods section, Driftsel’s threshold-FPR relationship did not follow the identity line. However, the most informative point is that Driftsel’s threshold-FPR relationship was inconsistent across population structures and scenarios, demonstrating a sensitivity of Driftsel to population structure. Under the island model population structure (Figure 4), the threshold-FPR curves were relatively shallow and consistent across different parameter combinations under this structure. In contrast, the stepping stones and hierarchical structures exhibited much steeper concave curves (Figure 4, middle and bottom panels). Notably, under the hierarchical structure, the curves varied substantially across scenarios with the same population structure. Under neutrality, the threshold-FPR relationship should remain stable regardless of the stabilizing selection strength (*ω*_*o*_), and differences between environments (Δ*θ*) being compared across panels, as these parameters affect only the selective scenarios and should not influence the null distribution.

These calibration results demonstrate that MONET (with log _*V*_ or the environment) is the only method among those tested that maintains reliable Type I error across diverse population structures, specifically including those with non-isotropic relatedness patterns.

### Statistical power

Under the island model (Figure 5), all three methods performed similarly across the range of selection scenarios tested, unsurprisingly given the fact that this population structure follows the *Q*_*ST*_ – *F*_*ST*_ assumption of isotropic relatedness. This demonstrates that MONET’s incorporation of population structure does not compromise power when the structure is isotropic. A notable distinction is how, for intermediate optima distances (Δ*θ* = 4.6) and weaker selection (*ω*_*o*_ = 22), the MONET method used with environmental information (referred to in the figures as “MONET w/environment”) has the highest power (TPR = 100%). The overall impact of including environmental covariates in the performance of MONET is further described below.

**Figure 5.**
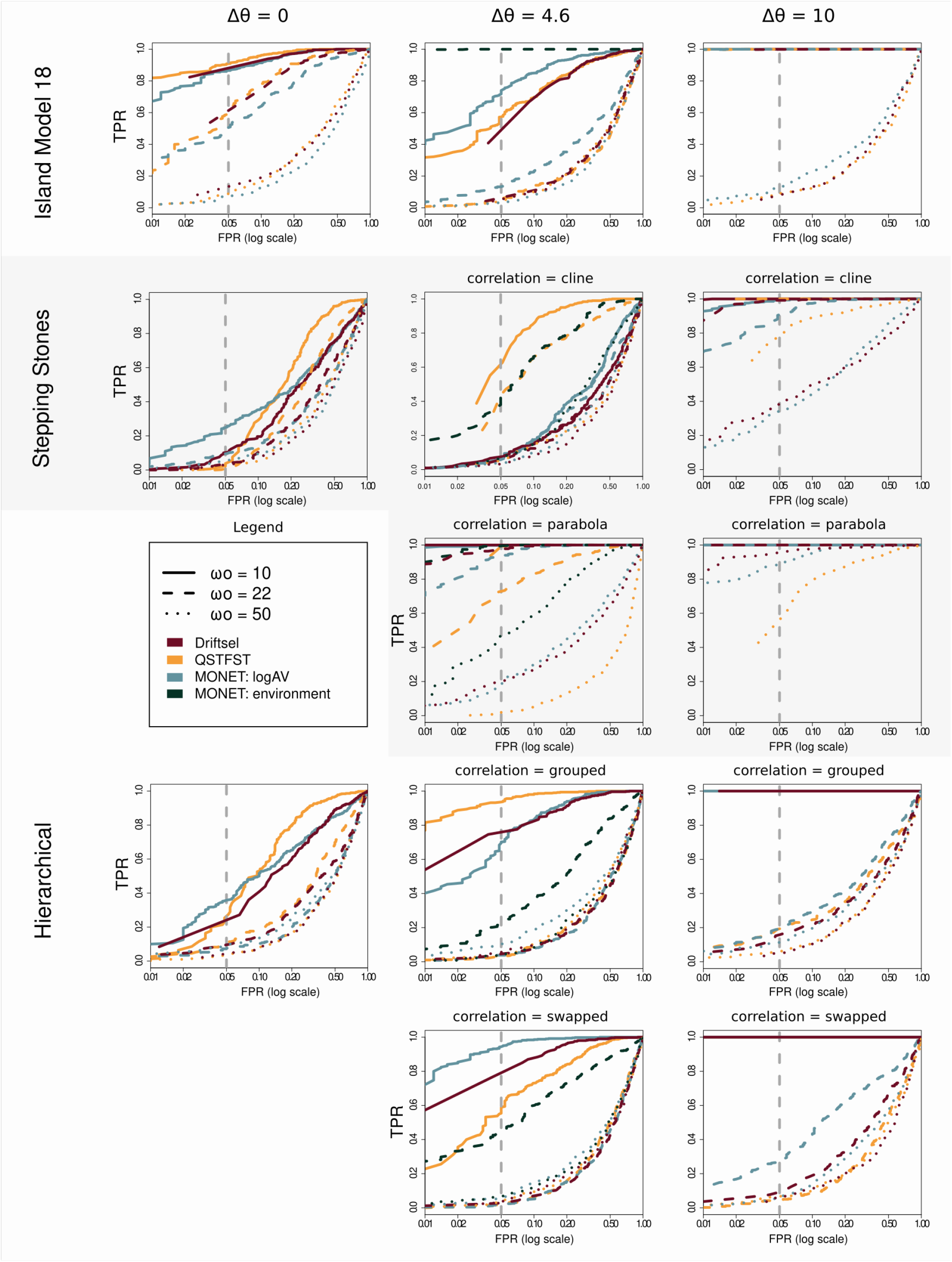
ROC graphs comparing true positive and false positive rates across the three methods for the three population structures with 18 subpopulations. The different graphs show different levels of environmental distance, with 0 indicating global adaptation. Within each graph, different colors describe different methods, and the different line types show different widths of the fitness optima.

Under the stepping stones structure (Figure 5), method performance diverged depending on the relationship between selective and neutral divergence patterns. Under global adaptation (Δ*θ* = 0), MONET maintained the highest power among all methods at FPR values near 0.05 (Table 2, and 5). Under local adaptation scenarios, the relationship between selective divergence and population structure critically affected method performance. When selective divergence followed a cline pattern that correlated with the stepping stones spatial arrangement (Δ*θ* = 4.6 and 10, cline), *Q*_*ST*_ – *F*_*ST*_ appeared to show higher power than structure-corrected methods (Table 2). However, when selective divergence followed a parabola pattern that was uncorrelated with neutral structure (Δ*θ* = 4.6 and 10, parabola), methods that account for population structure (MONET and Driftsel) substantially outperformed *Q*_*ST*_ – *F*_*ST*_ (Figure 5, middle panels). These differences are more pronounced under weak environmental differentiation (Δ*θ* = 4.6, and weaker selection strength *ω*_*o*_ = 22 or 50).

**Table 1.**
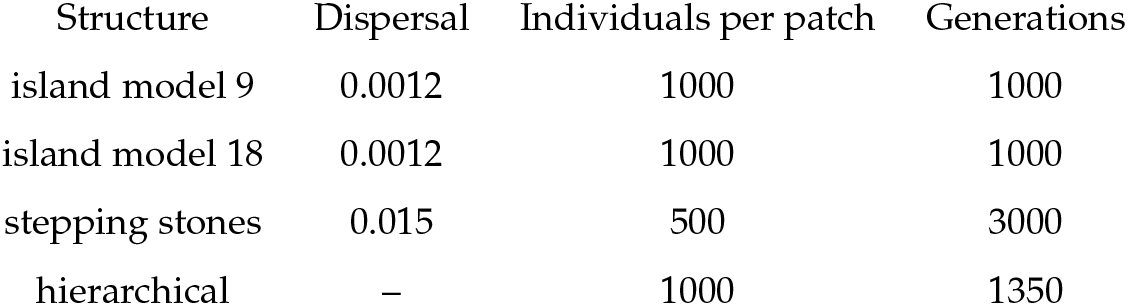
Description of main parameters of the simulated metapopulations.

| Structure | Dispersal | Individuals per patch | Generations |
| --- | --- | --- | --- |
| island model 9 | 0.0012 | 1000 | 1000 |
| island model 18 | 0.0012 | 1000 | 1000 |
| stepping stones | 0.015 | 500 | 3000 |
| hierarchical | – | 1000 | 1350 |

**Table 2.** True Positive Rate (TPR) at False Positive Rate (FPR) = 0.05 for different population structures and parameter combinations. Results are based on 500 replicates per scenario.

| Population structure | $\Delta\theta$ | $\omega_o$ | Correlation | QSTFST | MONET | Driftsel |
| --- | --- | --- | --- | --- | --- | --- |
| island model | 0 | 10 | — | 0.91 | 0.87 | 0.89 |
| island model | 0 | 22 | — | 0.60 | 0.51 | 0.57 |
| island model | 0 | 50 | — | 0.10 | 0.07 | 0.10 |
| island model | 10 | 10 | — | 1.00 | 1.00 | 1.00 |
| island model | 10 | 22 | — | 1.00 | 1.00 | 1.00 |
| island model | 10 | 50 | — | 0.09 | 0.13 | 0.09 |
| island model | 4.6 | 10 | — | 0.57 | 0.72 | 0.54 |
| island model | 4.6 | 22 | — | 0.06 | 0.13 | 0.05 |
| island model | 4.6 | 50 | — | 0.05 | 0.03 | 0.06 |
| stepping stones | 0.0 | 10 | — | 0.03 | 0.25 | 0.10 |
| stepping stones | 0.0 | 22 | — | 0.01 | 0.09 | 0.03 |
| stepping stones | 0.0 | 50 | — | 0.02 | 0.05 | 0.03 |
| stepping stones | 10.0 | 10 | cline | 1.00 | 0.99 | 1.00 |
| stepping stones | 10.0 | 10 | parabola | 1.00 | 1.00 | 1.00 |
| stepping stones | 10.0 | 22 | cline | 1.00 | 0.91 | 0.99 |
| stepping stones | 10.0 | 22 | parabola | 1.00 | 1.00 | 1.00 |
| stepping stones | 10.0 | 50 | cline | 0.78 | 0.34 | 0.39 |
| stepping stones | 10.0 | 50 | parabola | 0.56 | 0.89 | 0.96 |
| stepping stones | 4.6 | 10 | cline | 0.61 | 0.06 | 0.08 |
| stepping stones | 4.6 | 10 | parabola | 0.99 | 1.00 | 1.00 |
| stepping stones | 4.6 | 22 | cline | 0.44 | 0.06 | 0.06 |
| stepping stones | 4.6 | 22 | parabola | 0.73 | 0.94 | 0.98 |
| stepping stones | 4.6 | 50 | cline | 0.05 | 0.04 | 0.03 |
| stepping stones | 4.6 | 50 | parabola | 0.02 | 0.19 | 0.20 |
| hierarchical | 0 | 10 | — | 0.25 | 0.36 | 0.24 |
| hierarchical | 0 | 22 | — | 0.10 | 0.07 | 0.09 |
| hierarchical | 0 | 50 | — | 0.04 | 0.09 | 0.04 |
| hierarchical | 10 | 10 | grouped | 1.00 | 1.00 | 1.00 |
| hierarchical | 10 | 10 | swapped | 1.00 | 1.00 | 1.00 |
| hierarchical | 10 | 22 | grouped | 0.19 | 0.20 | 0.16 |
| hierarchical | 10 | 22 | swapped | 0.05 | 0.29 | 0.10 |
| hierarchical | 10 | 50 | grouped | 0.06 | 0.11 | 0.05 |
| hierarchical | 10 | 50 | swapped | 0.07 | 0.07 | 0.06 |
| hierarchical | 4.6 | 10 | grouped | 0.94 | 0.70 | 0.75 |
| hierarchical | 4.6 | 10 | swapped | 0.57 | 0.94 | 0.79 |
| hierarchical | 4.6 | 22 | grouped | 0.03 | 0.04 | 0.05 |
| hierarchical | 4.6 | 22 | swapped | 0.02 | 0.02 | 0.03 |
| hierarchical | 4.6 | 50 | grouped | 0.04 | 0.10 | 0.06 |
| hierarchical | 4.6 | 50 | swapped | 0.03 | 0.06 | 0.03 |

We observed similar results between the hierarchical structure and the stepping stones in terms of the power difference between methods. Under global adaptation (Δ*θ* = 0), MONET maintained the highest power under low false-positive rate thresholds. Under local adaptation, the alignment of selection patterns with population structure again proved critical. In the “grouped” scenario, where selective pressures aligned with hierarchical population groupings (Δ*θ* = 4.6, correlation = grouped), *Q*_*ST*_ – *F*_*ST*_ showed apparently higher power. However, in the “swapped” scenario where selection patterns were misaligned with population structure (Δ*θ* = 4.6, correlation = swapped), MONET substantially outperformed *Q*_*ST*_ – *F*_*ST*_ (Figure 5; Table 2). With Δ*θ* = 4.6 and *ω*_*o*_ = 10, MONET achieved TPR = 0.94 in the swapped scenario compared to 0.57 for *Q*_*ST*_ – *F*_*ST*_

When environmental information was incorporated into MONET’s framework (MONET w/environment), the method showed markedly improved power to detect selection compared to any of the standard approaches (Figure 5). MONET w/environment was even similarly or better performing than *Q*_*ST*_ – *F*_*ST*_ when the environment and population structure were strongly correlated.

### Effects of sampling design and common garden parameters

The performance of all methods varied with sampling design and study scale (Table 3). Increasing the number of sampled subpopulations from 9 to 18 improved power across all methods, with the greatest improvements observed in local adaptation scenarios (Δ*θ* = 4.6 and 10). This enhancement reflects increased information about the spatial pattern of phenotypic variation and population structure. The number of phenotyped individuals per population (F1 number) and breeding design also affected method performance, though impacts varied among population structures. For the island model and hierarchical structures, the two reduced breeding schemes (1 *×* 9 and 3 *×* 3) showed similar performance to the full breeding design. However, for the stepping stones structure, these effects were more pronounced. The 1 *×* 9 breeding design showed substantially better performance than the 3 *×* 3, with true positive rates approaching 0.94–1.00 versus 0.62 in the parabola scenario with Δ*θ* = 10 and *ω*_*o*_ = 22. The impact of reduced F1 sampling was strongest under stepping stones scenarios, where divergent selection correlated with neutral divergence (cline scenario), where fewer within-population samples led to reduced power to distinguish selection from neutral processes.

**Table 3.** True Positive Rate (TPR) at False Positive Rate (FPR) = 0.05 for different sampling schemes and population structures (Δ*θ* = 10, *ω*_*o*_ = 22). Each run represents a different spatial sampling strategy. Missing values on the table either indicate the method did not have values of TPR under the given FPR (Driftsel on 3×3 for stepping stones) or that the method did not accommodate the breeding scheme (e.g. *Q*_*ST*_ -*F*_*ST*_ on 1×9).

| Sampling or breeding scheme | Population structure | $Q_{ST}-F_{ST}$ | MONET | Driftsel |
| --- | --- | --- | --- | --- |
| Population number | island model | 0.45 | 0.45 | 0.43 |
| Population number | stepping stones cline | 0.18 | 0.10 | 0.11 |
| Population number | stepping stones parabola | 0.56 | 0.62 | 0.72 |
| Population number | hierarchical grouped | 0.37 | 0.35 | 0.33 |
| Population number | hierarchical swapped | 0.29 | 0.40 | 0.33 |
| F1 number | island model | 0.45 | 0.46 | 0.42 |
| F1 number | stepping stones cline | 0.16 | 0.05 | 0.06 |
| F1 number | stepping stones parabola | 0.02 | 0.98 | 0.99 |
| F1 number | hierarchical grouped | 0.37 | 0.35 | 0.33 |
| F1 number | hierarchical swapped | 0.29 | 0.40 | 0.33 |
| 3x3 | island model | 0.45 | 0.45 | 0.43 |
| 3x3 | stepping stones cline | 0.42 | 0.07 | — |
| 3x3 | stepping stones parabola | 0.90 | 0.94 | — |
| 3x3 | hierarchical grouped | 0.37 | 0.35 | 0.33 |
| 3x3 | hierarchical swapped | 0.29 | 0.40 | 0.33 |
| 1x9 | island model | — | 0.45 | 0.43 |
| 1x9 | stepping stones cline | — | 0.27 | 0.30 |
| 1x9 | stepping stones parabola | — | 1.00 | 1.00 |
| 1x9 | hierarchical grouped | — | 0.35 | 0.33 |
| 1x9 | hierarchical swapped | — | 0.40 | 0.33 |

### Results of the Quercus case study

In Figure 6, we show the metapopulation structure of the two Quercus metapopulations alongside representative **Θ**^***P***^ matrices from our simulated population structure.

**Figure 6.**
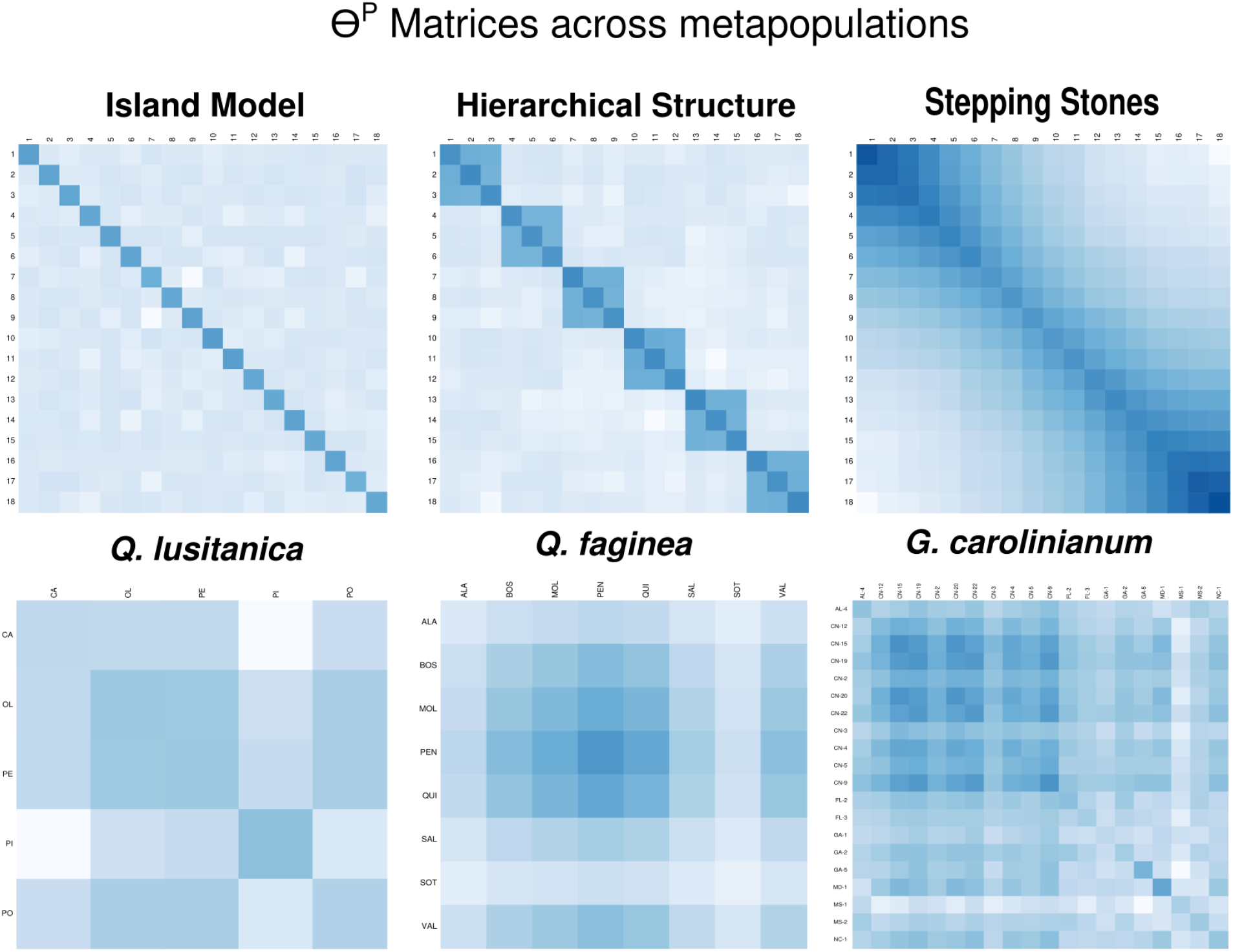
The **Θ**^***P***^ matrices of the different metapopulations discussed in this manuscript. We show the **Θ**^***P***^ matrix for one replicate of each of the simulated structures: island model, stepping stones, and hierarchical structures. And we show the population structure of the two metapopulations from the empirical case study in *Quercus*.

#### Testing for local or global adaptation

We first applied the base MONET model, which contains only the coancestry random effects, to test for signs of local or global adaptation through the log _*V*_ statistic (eq. 4). For *Q. faginea*, only specific leaf area in the well-watered treatment showed evidence of divergence beyond neutral expectation (log_*AV*_ median = 3.02, 95% CI [0.71, 6.21], *p* = 0.007). In *Q. lusitanica*, none showed significant log_*AV*_ . These results contrast with those obtained by Ramirez-Valiente *et al*. (2022) using *Q*_*ST*_ –*F*_*ST*_, who reported widespread evidence of selection on leaf morphology and physiology traits in *Q. faginea*, and a smaller subset of traits in *Q. lusitanica*.

#### Trait-climate association

The extended model adds PC_climate1_ and PC_climate2_ as fixed effects. We asked whether trait divergence tracks climate after accounting for neutral structure (eq. 6), testing for a more narrowly defined scenario of local adaptation, with more power. We found a similar species-level pattern to the original work by Ramirez-Valiente *et al*. (2022); *Q. faginea* traits show strong climate associations, and *Q. lusitanica* traits in general do not.

Specifically, in *Q. faginea*, MONET recovered all of the climate associations reported by Ramirez-Valiente *et al*. (2022), and identified three additional associations in physiological and growth traits, namely, we saw a significant association in absolute growth rate in the dry treatment, non-photochemical quenching in the dry treatment, and mass-based stomatal conductance in the well-watered treatment. In *Q. lusitanica*, MONET did not find the same results as the three Pearson associations reported by Ramirez-Valiente *et al*. (2022) (Area on PC1, mass-based stomatal conductance on PC1, and relative height growth rate on PC2). Unlike MONET, Pearson’s correlation on subpopulation means treats subpopulations as independent and ignores the non-independence arising from shared ancestry, which might explain the different results.

### Results of the Geranium case study

Out of the eleven traits in the original study’s trait list, seven showed a significant log_*AV*_ consistent with local adaptation: juvenile establishment (CI = [3.888, 11.255]), juvenile rosette size (CI = [1.108, 3.082]), adult rosette size (CI = [0.373,3.147]), height at harvest (CI = [0.602,2.719]), days to first flower (CI = [0.622, 2.477]), total aboveground biomass (CI = [0.616, 2.858]), and ten-seed weight (CI = [1.622, 3.217]). The contrast between countries, testing for divergence between the native and invasive ranges beyond neutral expectation, was significant for percentage of total nitrogen (*β* = +0.607, *p* = 5 *×* 10^−4^), days to first flower (*β* = +0.502, *p* = 0.007), and total aboveground biomass (*β* = − 0.555, *p* = 0.005).

Comparing these results to Shirk and Hamrick (2014) requires matching MONET’s two tests to their design of three *Q*_*ST*_ − *F*_*ST*_ tests. The contrast between countries corresponds to their interpretation of the comparison between the two countries (*Q*_*CT*_ − *F*_*CT*_), while the log_*AV*_ results would be related their within-country tests. However, we do not specifically test we did not test for local adaptation within the countries specifically to keep the case study simpler. The log_*AV*_ test here is showing how much of the divergence that cannot be explained by neutral processes is left after accounting for country contrast. It would would be possible to test this with MONET by including interaction between additive genetic effect and country effect. We did not include this test in order to keep the case study simpler. We show a full comparison between the two analyses in Table 4: For countries contrasts, 3 traits were significant with MONET and 2 with *Q*_*ST*_ –*F*_*ST*_, one (% total nitrogen) with similar results to both analyses. Within countries, 7 traits were significant with MONET, 5 in at least one country for *Q*_*ST*_ –*F*_*ST*_ . Of these, only 3 are have common results in both analysis.

**Table 4.** Comparison between Shirk and Hamrick (2014)’s three-way *Q*_*ST*_ –*F*_*ST*_ design and MONET. NS stands for No signal, + *QST* for significantly positive *Q*_*ST*_ –*F*_*ST*_ values which indicate divergent selection patters, and *Q*_*ST*_ for significantly negative *Q*_*ST*_ –*FST* indicating convergent selection patterns.

| Trait | Between countries | Country contrast - MONET | Within countries $Q_{SC}-F_{SC}$ | Local adaptation - MONET |
| --- | --- | --- | --- | --- |
| | $Q_{CT}-F_{CT}$ | | | |
| Juvenile establishment time | NS | NS | + $Q_{ST}$ (China) | local adaptation |
| Juvenile rosette size | + $Q_{ST}$ | NS | + $Q_{ST}$ (both) | local adaptation |
| Specific leaf area | NS | NS | + $Q_{ST}$ (China) | NS |
| Adult rosette size | NS | NS | NS | local adaptation |
| Percentage of total nitrogen | + $Q_{ST}$ | + $Q_{ST}$ | + $Q_{ST}$ (China); (US) | NS |
| $\delta^{13}C$ | NS | NS | NS | NS |
| Height | NS | NS | NS | local adaptation |
| Days to first flower | NS | + $Q_{ST}$ | NS | local adaptation |
| Reproductive ratio | NS | NS | - $Q_{ST}$ (China) | NS |
| Aboveground biomass | NS | + $Q_{ST}$ | + $Q_{ST}$ (both countries) | local adaptation |
| Ten-seed weight | NS | NS | NS | local adaptation |

## Discussion

The aim of this work was to present a flexible and robust neutral model for trait evolution that can be used to implement hypothesis tests including tests for trait association with environmental variables, local and global adaptation tests, and the influence of other factors on trait distribution within a metapopulation. We do so by presenting how MONET expands the method proposed by (do O *et al*. 2025), then show via extensive simulations that only MONET maintained calibration even when the population structure became non-isotropic. Finally, the two empirical data sets reanalyses that we carried out illustrate the practical consequences of a flexible and robust neutral model that considers population structure.

After introducing our framework, we assess whether currently available methods for detecting selection on quantitative traits remain reliable across population structures. The findings of MONET’s consistent calibration have important practical implications, as natural populations rarely exhibit the isotropic gene flow, equal ancestral contributions, and similar effective population sizes assumed by the island model. Importantly, MONET remains well-calibrated while maintaining similar or greater power than the other methods, except for *Q*_*ST*_ –*F*_*ST*_ when local adaptation and population structure are confounded, which we discuss further below. Proper calibration is an essential feature of a statistical test. Here, it allows for a more reliable inference about adaptive divergence in a metapopulation.

The miscalibration of *Q*_*ST*_ – *F*_*ST*_ under non-isotropic structure has been recognized as a theoretical concern (do O *et al*. 2025; de Villemereuil *et al*. 2022; Ovaskainen *et al*. 2011), and our simulation results quantify the practical magnitude of this issue. The consistent inflation of false positive rates under non-isotropic structure means that researchers using *Q*_*ST*_ – *F*_*ST*_ in spatially structured systems may incorrectly infer selection when phenotypic divergence may be simply reflecting neutral processes shaped by demographic history. The consistent inflation of false positive rates in *Q*_*ST*_ – *F*_*ST*_ emphasizes the need for a more complete neutral model that accounts for the topology of metapopulations, such as MONET. Similarly, our results suggest Driftsel, even while not expected to be calibrated, may be unreliable as the behaviour of the threshold to FPR relationship was sensitive to population structure and selection parameters.

Methods that fail to account for population structure will likely confound neutral differentiation with adaptive divergence when these patterns happen to align. Our results clearly illustrate this issue: *Q*_*ST*_ – *F*_*ST*_ often appeared to have higher power than structure-corrected methods when environmental differences aligned with population structure, such as in the stepping stones cline scenario and hierarchical grouped scenario (Table 2). However, this apparent advantage is misleading. Because the neutral model underlying the *Q*_*ST*_ – *F*_*ST*_ approach does not accommodate non-isotropic population structure, it conflates neutral differentiation arising from demographic history with adaptive divergence, which also increases the false positive rate.

Many real-world populations show spatial structure arising from isolation-by-distance, discrete geographic barriers, or hierarchical patterns of colonization and divergence. For instance, Viricel and Rosel (2014) documented hierarchical genetic structure in Atlantic spotted dolphins (*Stenella frontalis*), where the highest level of divergence separated oceanic and shelf ecotypes, with further subdivision within the continental shelf cluster into three distinct groups. This complex structure can lead to a high false-positive rate if the purpose of the study is to identify quantitative trait divergence using currently popular methods. Similarly, Pisa *et al*. (2015) identified a hierarchical structure in Fire Salamander (*Salamandra salamandra*) populations in Northern Italy, where populations were separated into two groups, with one of them showing further subdivision into four clusters. Isolation-by-distance patterns, similar to the stepping stones structure, also seem to be common. For example, Cassel-Lundhagen *et al*. (2013) found that the northern pine processionary moth (*Thaumetopoea pinivora*) exhibited genetic structure consistent with isolation-by-distance across its scattered European range, while Gold *et al*. (2001) documented that red drum (*Sciaenops ocellatus*) populations in the Gulf of Mexico follow a stepping-stone model where gene flow decreases continuously with geographic distance. Maliti *et al*. (2014) showed that *Anopheles arabiensis* exhibited stepping-stone structure with extensive but geographically restricted gene flow (average *F*_*ST*_ = 0.015) and significant isolation-by-distance. In all such cases, the neutral model underlying MONET would provide a faithful null against which to evaluate trait divergence, since it accommodates these heterogeneous relatedness structures.

Another important factor is the variation in effective sizes between subpopulations, which we did not directly test in this work. MONET accommodates this through the estimation of population-specific components, more precisely in the diagonal elements of **Θ**^*p*^. Differences in effective population sizes and demographic histories between subpopulations create heterogeneity in within-population relatedness. This is particularly relevant for conservation scenarios involving small isolated populations and larger source populations. For instance, Khan *et al*. (2021) documented substantial differences in genomic inbreeding between a small isolated tiger population (mean *F*_*ROH*_ = 0.57) and two large connected populations (mean *F*_*ROH*_ = 0.35 and 0.46), reflecting their distinct demographic histories and levels of connectivity. Such variation in inbreeding and effective population size directly translates into heterogeneous relatedness structures that violate the assumptions of *Q*_*ST*_ –*F*_*ST*_ comparison.

Our reanalysis of the Mediterranean Quercus dataset from Ramirez-Valiente *et al*. (2022) provides an empirical example of the issues we presented. In *Q. faginea*, MONET detected only “specific leaf area” in the well-watered treatment, showing clear signs of local adaptation, and in *Q. lusitanica* MONET did not detect convincing evidence that quantitative trait divergence exceeded neutral expectations, despite the widespread signal inferred from *Q*_*ST*_ –*F*_*ST*_ . Notably, Ramirez-Valiente *et al*. (2022) themselves observed discrepancies between *Q*_*ST*_ –*F*_*ST*_ results and Driftsel, with *Q*_*ST*_ –*F*_*ST*_ flagging more traits as under selection. In these results, *Q*_*ST*_ –*F*_*ST*_ detected the largest number of traits under selection, followed by Driftsel, and finally MONET, which detected the fewest.

The reanalyzing of the work on Geranium by Shirk and Hamrick (2014) allowed us to resolve a problem they posed in their discussion, which was the potential lack of statistical power in their “*Q*_*CT*_ –*F*_*CT*_ “ analyses with only two subpopulations. By including country as a fixed effect in MONET, we could address the question of whether there was divergent selection between the invasive and the native subpopulations without decreasing the number of samples, or misrepresenting the number of populations, both of which result in a loss of power. While we, similarly to the original results, found that invasive Chinese individuals had significantly higher nitrogen in their leaves, we also found that Chinese plants flowered significantly later and had significantly smaller total aboveground biomass, which the original results did not find, possibly related to the power issue mentioned just above. These two new significant results follow what Shirk and Hamrick (2014) expected to find in relation to the new, harsher climatic conditions in China. The authors of the original work found a pattern of smaller plants flowering later in the season, which would be expected as a result of adaptation to larger seasonal variations experienced by the Chinese subpopulations.

While in one of the reanalyses (Quercus) we found fewer traits under divergence selection, and in the second (Geranium) we found more than the original results, we argue that both are consequences of a neutral model that more robustly represents and leverages population structure into hypothesis testing. Although Ramirez-Valiente *et al*. (2022) reports weak population structure in both Quercus species, in both of them, pairwise *F*_*ST*_ already showed a wide variation of over an order of magnitude differences (range = 0.002 to 0.073, and 0.006 - 0.089). This pattern reported by the original authors already indicates a non-isotropic structure, even if weak on average. Thus, we interpret MONET’s fewer detections as consistent with appropriate calibration rather than necessarily reduced sensitivity. In the Geranium dataset, by contrast, an important limiting factor in their work was the loss of statistical power from collapsing twenty subpopulations into two, as noted by Shirk and Hamrick (2014). By incorporating country contrast as a fixed effect within a single model, MONET recovers additional signals without misrepresenting population structure. We can only infer more general confidence in MONET on the basis of our simulation analysis. Thus, in both case studies, as in any empirical data analyzed, we cannot firmly conclude on the confidence in each particular result compared to the original studies.

Traditional *Q*_*ST*_ – *F*_*ST*_ comparisons provide an overall test of whether divergence exceeds neutral expectations under an island model, but cannot identify which environmental factors drive adaptation. Researchers have typically addressed this limitation by conducting separate analyses: first testing for local adaptation using the *Q*_*ST*_ – *F*_*ST*_ method, then separately regressing population mean trait values against environmental variables to identify putative selective agents (e.g. Marin *et al*. 2020; Cooper *et al*. 2021). This two-step approach has two drawbacks: *(i)* it does not leverage environmental information to increase power for detecting local adaptation; and *(ii)* the regression steps suffer from an inflated error rate because it does not account for non-independence between populations due to neutral processes. MONET’s mixed-effects framework naturally accommodates environmental covariates while accounting for population structure.

Notably, Karhunen *et al*. (2014) extended the Driftsel framework with an “H”-test. This statistic asks whether populations in similar habitats are more phenotypically similar than expected from their shared ancestry and genetic correlations alone. It incorporates habitat information directly into the neutrality test for quantitative traits. MONET differs from Driftsel’s H-test in how environmental information is used. Rather than comparing habitat and trait distance matrices after fitting the model, MONET estimates environmental effects as fixed-effect coefficients within the mixed model itself, together with the neutral structure. It produces the environment association slope and the log_*AV*_ value through the same fit. These differences impact the interpretation of the results, being much more intuitive when using MONET.

An important design choice in MONET is to keep the between-population and the within-population coancestry structures separate, rather than collapsing them into a single relatedness matrix. Although the unified formulation is possible, it would tend to absorb any unexplained signal into a single variance component, including signals arising from local adaptation to environmental drivers that are not explicitly modeled.

By explicitly modeling how environmental variation predicts phenotypic divergence, MONET with environmental covariates can more effectively partition variance between selective and neutral processes. This approach also provides an additional insight into the ecological drivers of divergence, allowing researchers to test specific hypotheses about which environmental factors are most important for adaptation. This approach being correlative, it requires that the tested variable is a good proxy for the causal factor(s) of local adaptation, and cannot in itself distinguish between a proxy and the causal variable. It also requires a linear relationship between the environmental variable and phenotypic optima in the simplest form of the model. Fortunately, MONET’s flexibility allows researchers to model more complex relationships by incorporating non-linear terms in the fixed effects formula. For instance, quadratic or polynomial terms can capture curvilinear relationships, and categorical environmental variables can accommodate discrete habitat types with distinct selective pressures. Researchers should specify these more complex relationships based on *a priori* biological hypotheses about how environmental factors influence selective pressures.

More broadly, when including environmental covariates, MONET can be seen as the quantitative-genetic analogue of Genotype–Environment Association (GEA) studies. GEA methods test for associations between allele frequencies and environmental variables while accounting for neutral population structure (Coop *et al*. 2010; Frichot *et al*. 2013), while MONET tests for the analogous association at the level of the phenotype using the coancestry matrices to control for that same neutral structure. It can therefore be incorporated into integrated pipelines for the study of local adaptation, such as that of de Villemereuil *et al*. (2018), contributing to the environment–phenotype link within the broader environment–phenotype–genotype chain of associations that together characterize local adaptation.

A last strength of this environmental approach is that MONET can test several variables in a multivariate framework using multiple regression, although this was also something provided by Driftsel (Karhunen *et al*. 2014) when using the H-statistics. The substantial power gain achieved by incorporating environmental covariates into MONET highlights the value of collecting environmental data alongside phenotypic and genetic information in studies of adaptive divergence.

Another advantage of MONET being based on a mixed modeling framework is that when traits are non-Gaussian (e.g., binary, count, or bounded data), it can accommodate them using generalized linear mixed models (GLMMs) by choosing an appropriate family and link function in `brms`. In such cases, inference occurs on the latent scale (the scale of the linear predictor), which is standard practice in quantitative genetics for non-normal traits and should not impact MONET’s testing procedures (both using log_*AV*_ and the environmental test). Note that back-transformation of the variances (De Villemereuil *et al*. 2016) is not needed to compute log_*AV*_, as the transformation factor would cancel out in the computation of the ratio.

The different applications shown in this work point to the broad role of MONET as a neutral model of trait evolution. Although the idea of neutral models of quantitative-trait evolution has been suggested before, it has not yet been developed to accommodate the diversity of natural populations. MONET relaxes the assumptions on population structure, providing a general neutral framework for phenotypic traits across a wide range of life histories, breeding designs, trait distributions, and demographic structures.

## Data availability

Scripts and data supporting this study are available at: https://github.com/isadoo/Testing_MONET and the package can be found at: https://github.com/isadoo/MONET.

## Supporting Information

### Description of tested methods

*Q*_*ST*_ ***–*** *F*_*ST*_ Whitlock and Guillaume (2009) approach follows the same premise as the classical *Q*_*ST*_ – *F*_*ST*_ comparison proposed by Spitze (1993) in which under neutral evolution, quantitative trait differentiation (*Q*_*ST*_) should match neutral marker differentiation (*F*_*ST*_). Deviations from this expectation provide evidence of selection. The issue brought to light by Whitlock (2008) was that in *Q*_*ST*_–*F*_*ST*_ comparisons, both statistics are subject to sampling error, and their distributions depend on demographic parameters. Thus, instead of comparing point estimates, *Q*_*ST*_ must be compared to the expected distribution of neutral *Q*_*ST*_ values given the observed *F*_*ST*_.

Throughout this manuscript, we will refer to *Q*_*ST*_ – *F*_*ST*_ as the parametric simulation approach developed by Whitlock and Guillaume (2009). Their approach is based on building null distributions through a parametric bootstrap. To incorporate uncertainty, *F*_*ST*_ should be measured for each simulation replicate by resampling neutral loci with replacement. For estimating *Q*_*ST*_, *V*_*W*_ and *V*_*B*_ will be calculated using different approaches. *V*_*W*_ is estimated based on the breeding design, with the sampling distribution being simulated using a parametric bootstrap assuming the mean squares follow a *χ*^2^ distribution (Lewontin and Krakauer 1973). For *V*_*B*_, rather than using the observed variance, the method calculates the expected *V*_*B*_ under neutrality, given the observed *F*_*ST*_ and *V*_*W*_ . Note that this test assumes that trait means between subpopulations are normally distributed, and that *V*_*W*_ is equal across all subpopulations.

#### Driftsel

Driftsel (Ovaskainen *et al*. 2011; Karhunen *et al*. 2013) addresses issues related to the isotropic assumption of *Q*_*ST*_–*F*_*ST*_ methods. Ovaskainen *et al*. (2011) approach is based on the idea that under neutrality the covariance in mean trait value between two populations (*X* and *Y*) should be proportional to their coancestry: 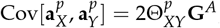, where **G**^*A*^ is the ancestral additive genetic variance-covariance matrix.

Driftsel’s test for evidence of selection uses the *S*-statistic. Driftsel computes the probability density of observing the actual configuration of population effects **a**^*p*^ under the multivariate normal distribution expected from drift: 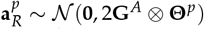. The *S*-statistic is then defined based on where this observed probability density falls relative to the distribution of densities expected from random realizations of the neutral process, as shown in equation 7:

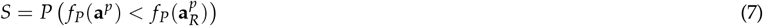

Values of *S* close to 0 or 1 (in the tails of the expected distribution) provide evidence against neutrality, while values near 0.5 are consistent with drift. Here, we follow the authors’ suggestions in (Ovaskainen *et al*. 2011; Karhunen *et al*. 2013) and use values of *S* lower than 0.2 to indicate global adaptation and values higher than 0.8 to indicate local adaptation.

One issue with the *S*-statistic is that it does not have an expected sampling distribution to which it can be compared. Indeed, Driftsel measures *S* through a Bayesian method that generates posterior samples. In our study, we get one posterior sample of *s* for each independent replicate of the scenario we are analyzing. The value of *S* we use is then the average of this posterior sample. Thus, because of the central limit theorem, the sampling distribution of *S* will be asymptotically Gaussian. However, we do not have a clear expectation for the standard deviation of this sampling distribution.

**Table S.1.**
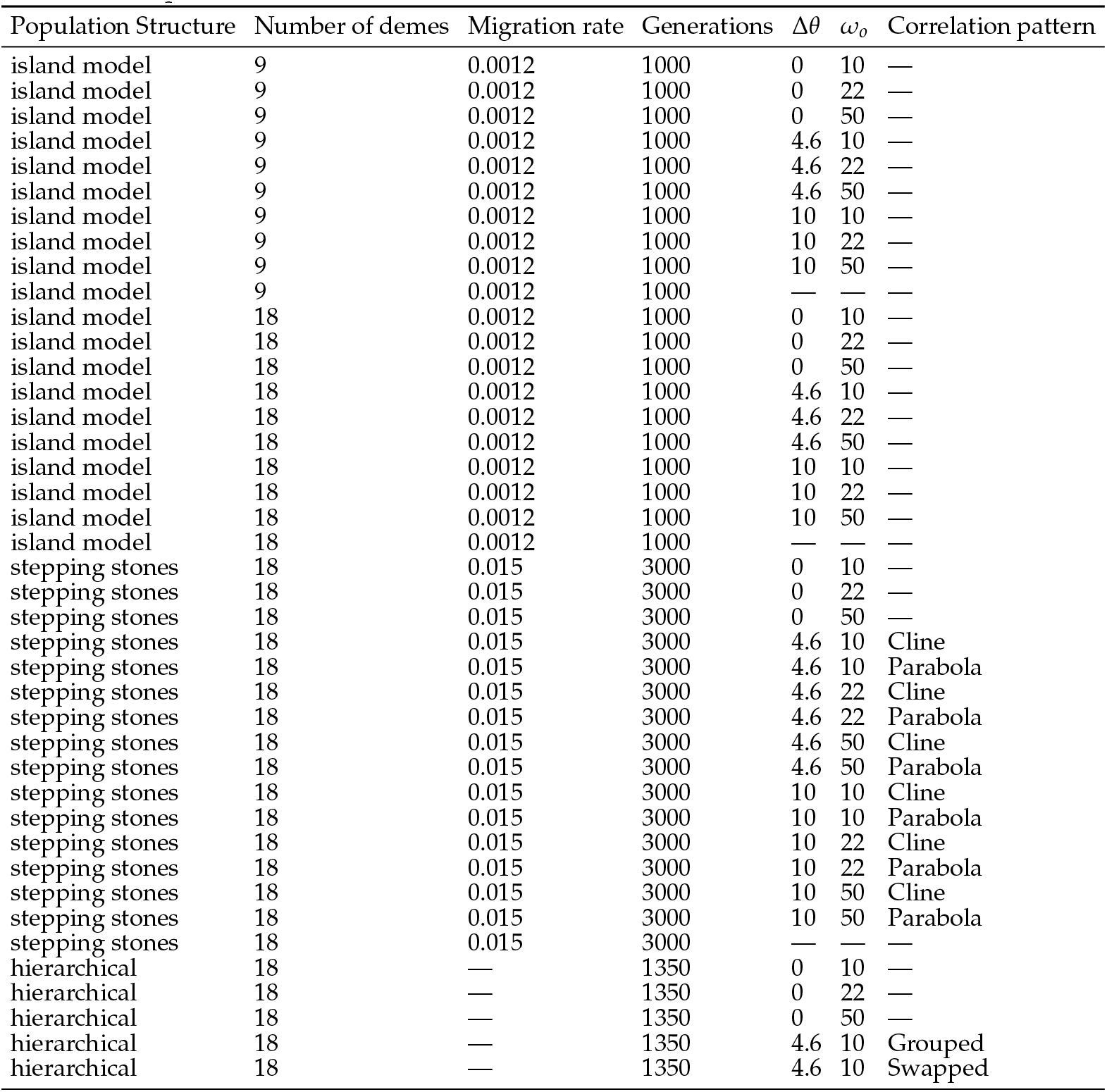
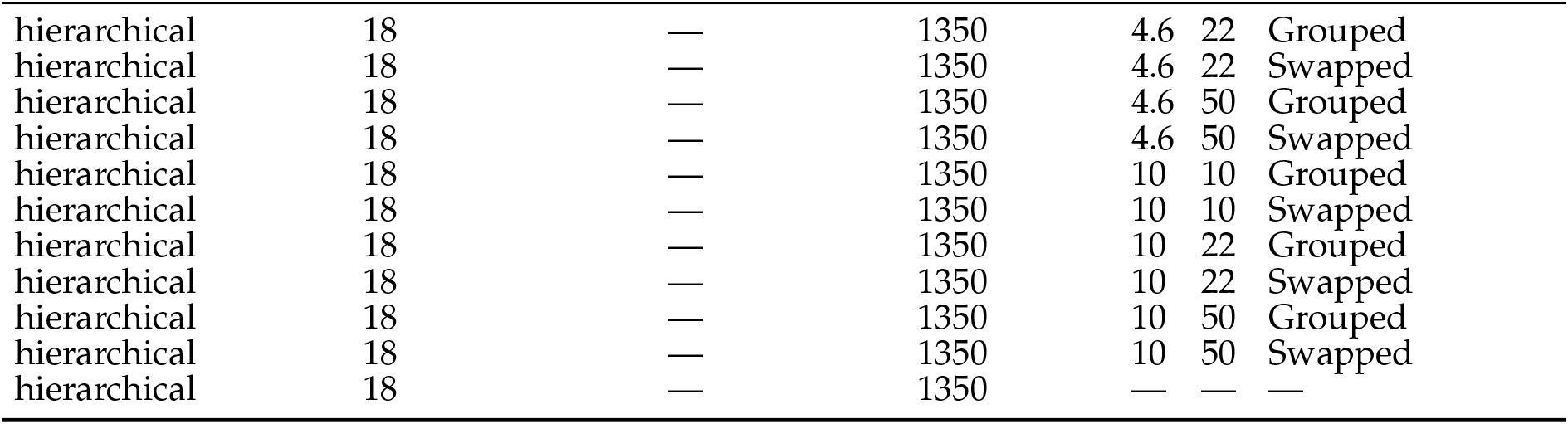
Description of the main parameters of the different scenarios simulated.

| Population Structure | Number of demes | Migration rate | Generations | $\Delta\theta$ | $\omega_o$ | Correlation pattern |
| --- | --- | --- | --- | --- | --- | --- |
| island model | 9 | 0.0012 | 1000 | 0 | 10 | — |
| island model | 9 | 0.0012 | 1000 | 0 | 22 | — |
| island model | 9 | 0.0012 | 1000 | 0 | 50 | — |
| island model | 9 | 0.0012 | 1000 | 4.6 | 10 | — |
| island model | 9 | 0.0012 | 1000 | 4.6 | 22 | — |
| island model | 9 | 0.0012 | 1000 | 4.6 | 50 | — |
| island model | 9 | 0.0012 | 1000 | 10 | 10 | — |
| island model | 9 | 0.0012 | 1000 | 10 | 22 | — |
| island model | 9 | 0.0012 | 1000 | 10 | 50 | — |
| island model | 9 | 0.0012 | 1000 | — | — | — |
| island model | 18 | 0.0012 | 1000 | 0 | 10 | — |
| island model | 18 | 0.0012 | 1000 | 0 | 22 | — |
| island model | 18 | 0.0012 | 1000 | 0 | 50 | — |
| island model | 18 | 0.0012 | 1000 | 4.6 | 10 | — |
| island model | 18 | 0.0012 | 1000 | 4.6 | 22 | — |
| island model | 18 | 0.0012 | 1000 | 4.6 | 50 | — |
| island model | 18 | 0.0012 | 1000 | 10 | 10 | — |
| island model | 18 | 0.0012 | 1000 | 10 | 22 | — |
| island model | 18 | 0.0012 | 1000 | 10 | 50 | — |
| island model | 18 | 0.0012 | 1000 | — | — | — |
| stepping stones | 18 | 0.015 | 3000 | 0 | 10 | — |
| stepping stones | 18 | 0.015 | 3000 | 0 | 22 | — |
| stepping stones | 18 | 0.015 | 3000 | 0 | 50 | — |
| stepping stones | 18 | 0.015 | 3000 | 4.6 | 10 | Cline |
| stepping stones | 18 | 0.015 | 3000 | 4.6 | 10 | Parabola |
| stepping stones | 18 | 0.015 | 3000 | 4.6 | 22 | Cline |
| stepping stones | 18 | 0.015 | 3000 | 4.6 | 22 | Parabola |
| stepping stones | 18 | 0.015 | 3000 | 4.6 | 50 | Cline |
| stepping stones | 18 | 0.015 | 3000 | 4.6 | 50 | Parabola |
| stepping stones | 18 | 0.015 | 3000 | 10 | 10 | Cline |
| stepping stones | 18 | 0.015 | 3000 | 10 | 10 | Parabola |
| stepping stones | 18 | 0.015 | 3000 | 10 | 22 | Cline |
| stepping stones | 18 | 0.015 | 3000 | 10 | 22 | Parabola |
| stepping stones | 18 | 0.015 | 3000 | 10 | 50 | Cline |
| stepping stones | 18 | 0.015 | 3000 | 10 | 50 | Parabola |
| stepping stones | 18 | 0.015 | 3000 | — | — | — |
| hierarchical | 18 | — | 1350 | 0 | 10 | — |
| hierarchical | 18 | — | 1350 | 0 | 22 | — |
| hierarchical | 18 | — | 1350 | 0 | 50 | — |
| hierarchical | 18 | — | 1350 | 4.6 | 10 | Grouped |
| hierarchical | 18 | — | 1350 | 4.6 | 10 | Swapped |
| hierarchical | 18 | — | 1350 | 4.6 | 22 | Grouped |
| hierarchical | 18 | — | 1350 | 4.6 | 22 | Swapped |
| hierarchical | 18 | — | 1350 | 4.6 | 50 | Grouped |
| hierarchical | 18 | — | 1350 | 4.6 | 50 | Swapped |
| hierarchical | 18 | — | 1350 | 10 | 10 | Grouped |
| hierarchical | 18 | — | 1350 | 10 | 10 | Swapped |
| hierarchical | 18 | — | 1350 | 10 | 22 | Grouped |
| hierarchical | 18 | — | 1350 | 10 | 22 | Swapped |
| hierarchical | 18 | — | 1350 | 10 | 50 | Grouped |
| hierarchical | 18 | — | 1350 | 10 | 50 | Swapped |
| hierarchical | 18 | — | 1350 | — | — | — |

## Literature cited

Beavis W, Espinosa K, Newell M, Mahama AA. 2023. Quantitative Genetics for Plant Breeding. Iowa State University Digital Press. Ames, Iowa. suza_quantitative_2023.

Bürkner PC. 2018. Advanced Bayesian Multilevel Modeling with the R Package brms. The R Journal. 10:395–411.

Cassel-Lundhagen A, Ronnås C, Battisti A, Wallén J, Larsson S. 2013. Stepping-stone expansion and habitat loss explain a peculiar genetic structure and distribution of a forest insect. Molecular Ecology. 22:3362–3375.

Cockerham CC, Tachida H. 1987. Evolution and maintenance of quantitative genetic variation by mutations. Proceedings of the National Academy of Sciences. 84:6205–6209.

Coop G, Witonsky D, Di Rienzo A, Pritchard JK. 2010. Using environmental correlations to identify loci underlying local adaptation. Genetics. 185:1411–1423.

Cooper H, Allan G, Andrews L, Best R, Grady K et al. 2021. Climate-driven divergent selection in a foundation tree species: QST-FST evidence from multiple common gardens. Authorea Preprints. .

de Villemereuil P, Gaggiotti OE, Goudet J. 2022. Common garden experiments to study local adaptation need to account for population structure. Journal of Ecology. 110:1005–1009.

de Villemereuil P, Mouterde M, Gaggiotti OE, Till-Bottraud I. 2018. Patterns of phenotypic plasticity and local adaptation in the wide elevation range of the alpine plant Arabis alpina. Journal of Ecology. 106:1952–1971.

De Villemereuil P, Schielzeth H, Nakagawa S, Morrissey M. 2016. General methods for evolutionary quantitative genetic inference from generalized mixed models. Genetics. 204:1281–1294.

do O I, Gaggiotti OE, de Villemereuil P, Goudet J. 2025. A method for identifying local adaptation in structured populations. PLoS Genetics. 21:e1011871.

do O I, Gaggiotti OE, de Villemereuil P, Goudet J. 2026. MONET: Model of Neutral Evolution of Traits.

Fawcett T. 2004. ROC graphs: Notes and practical considerations for researchers. Machine learning. 31:1–38.

Fick SE, Hijmans RJ. 2017. WorldClim 2: new 1-km spatial resolution climate surfaces for global land areas. International journal of climatology. 37:4302–4315.

Frichot E, Schoville SD, Bouchard G, François O. 2013. Testing for Associations between Loci and Environmental Gradients Using Latent Factor Mixed Models. Molecular Biology and Evolution. 30:1687–1699.

Gold J, Burridge C, Turner T. 2001. A modified stepping-stone model of population structure in red drum, Sciaenops ocellatus (Sciaenidae), from the northern Gulf of Mexico. Genetica. 111:305–317.

Hargreaves GH, Samani ZA. 1985. Reference crop evapotranspiration from temperature. Applied engineering in agriculture. 1:96–99.

Huisman J. 2017. Pedigree reconstruction from SNP data: parent-age assignment, sibship clustering and beyond. Molecular ecology resources. 17:1009–1024.

Jones OR, Wang J. 2010. COLONY: a program for parentage and sibship inference from multilocus genotype data. Molecular ecology resources. 10:551–555.

Karhunen M, Merilä J, Leinonen T, Cano J, Ovaskainen O. 2013. driftsel: an R package for detecting signals of natural selection in quantitative traits. Molecular Ecology Resources. 13:746– 754.

Karhunen M, Ovaskainen O, Herczeg G, Merilä J. 2014. Bringing habitat information into statistical tests of local adaptation in quantitative traits: a case study of nine-spined sticklebacks. Evolution; international journal of organic evolution. 68:559–568.

Khan A, Patel K, Shukla H, Viswanathan A, van der Valk T et al. 2021. Genomic evidence for inbreeding depression and purging of deleterious genetic variation in Indian tigers. Proceedings of the National Academy of Sciences. 118:e2023018118.

Kimura M. 1983. Rare variant alleles in the light of the neutral theory. Molecular biology and evolution. 1:84–93.

Kimura M, others. 1968. Evolutionary rate at the molecular level. Nature. 217:624–626.

Kimura M, Weiss GH. 1964. The stepping stone model of population structure and the decrease of genetic correlation with distance. Genetics. 49:561.

Lande R. 1992. Neutral theory of quantitative genetic variance in an island model with local extinction and colonization. Evolution; international journal of organic evolution. 46:381–389.

Lewontin RC, Krakauer J. 1973. Distribution of gene frequency as a test of the theory of the selective neutrality of polymorphisms. Genetics. 74:175–195.

Lynch M. 1994. Neutral models of phenotypic evolution, In: Real LA, editor, Ecological genetics, Princeton University Press. pp. 86–108. Section: 5.

Lynch M, Walsh B, others. 1998. Genetics and analysis of quantitative traits. volume 1. Sinauer Sunderland, MA.

Maliti D, Ranson H, Magesa S, Kisinza W, Mcha J et al. 2014. Islands and stepping-stones: comparative population structure of Anopheles gambiae sensu stricto and Anopheles arabiensis in Tanzania and implications for the spread of insecticide resistance. PLoS One. 9:e110910.

Marin S, Gibert A, Archambeau J, Bonhomme V, Lascoste M et al. 2020. Potential adaptive divergence between subspecies and populations of snapdragon plants inferred from QST–FST comparisons. Molecular Ecology. 29:3010–3021.

Neuenschwander S, Michaud F, Goudet J. 2019. QuantiNemo 2: a Swiss knife to simulate complex demographic and genetic scenarios, forward and backward in time. Bioinformatics (Oxford, England). 35:886–888.

Ovaskainen O, Karhunen M, Zheng C, Arias JMC, Merilä J. 2011. A new method to uncover signatures of divergent and stabilizing selection in quantitative traits. Genetics. 189:621–632.

Pisa G, Orioli V, Spilotros G, Fabbri E, Randi E et al. 2015. Detecting a hierarchical genetic population structure: the case study of the Fire Salamander (Salamandra salamandra) in Northern Italy. Ecology and Evolution. 5:743–758.

Poggio L, De Sousa LM, Batjes NH, Heuvelink G, Kempen B et al. 2021. SoilGrids 2.0: producing soil information for the globe with quantified spatial uncertainty. Soil. 7:217–240.

Ramirez-Valiente JA, Sole-Medina A, Robledo-Arnuncio JJ, Ortego J. 2022. Genomic data and common garden experiments reveal climate-driven selection on ecophysiological traits in two Mediterranean oaks. Molecular ecology. .

Shi H, Yin G. 2021. Reconnecting p-value and posterior probability under one-and two-sided tests. The American Statistician. 75:265–275.

Shirk RY, Hamrick JL. 2014. Multivariate adaptation but no increase in competitive ability in invasive Geranium carolinianum L.(Geraniaceae). Evolution; international journal of organic evolution. 68:2945–2959.

Slatkin M, Voelm L. 1991. FST in a hierarchical island model. Genetics. 127:627–629.

Spitze K. 1993. Population structure in Daphnia obtusa: quantitative genetic and allozymic variation. Genetics. 135:367–374.

Trabucco A, Zomer RJ. 2010. Global soil water balance geospatial database. CGIAR consortium for spatial information. .

Viricel A, Rosel PE. 2014. Hierarchical population structure and habitat differences in a highly mobile marine species: the Atlantic spotted dolphin. Molecular ecology. 23:5018–5035.

Watterson G. 1975. On the number of segregating sites in genetical models without recombination. Theoretical population biology. 7:256–276.

Whitlock MC. 1999. Neutral additive genetic variance in a metapopulation. Genetics Research. 74:215–221.

Whitlock MC. 2008. Evolutionary inference from QST. Molecular ecology. 17:1885–1896.

Whitlock MC, Guillaume F. 2009. Testing for Spatially Divergent Selection: Comparing QST to FST. Genetics. 183:1055–1063.

Wright S. 1949. The genetical structure of populations. Annals of eugenics. 15:323–354.

Yeaman S. 2022. Evolution of polygenic traits under global vs local adaptation. Genetics. 220:iyab134. tex.eprint: https://academic.oup.com/genetics/articlepdf/220/1/iyab134/42074616/iyab134.pdf.

